# A thirty-year trend of increasing clinical orientation at the National Institutes of Health

**DOI:** 10.64898/2025.12.16.694423

**Authors:** Brad L. Busse, James M. Tucker, Summer E. Allen, George M. Santangelo, Kristine A. Willis

**Author notes:** Woodley Park Institute, Washington, DC, 20008.

## Abstract

It is widely recognized that funding for biomedical research supports the development of major medical advances. However, little systematic effort has been made to determine whether a link exists between the types of funding opportunities that are available to scientists and progress towards new treatments in the clinic. To better understand this relationship, we analyzed the funding opportunities offered by the National Institutes of Health (NIH) over a span of thirty-two years, together with the resulting portfolio of applications and awards. We found NIH funding opportunity announcements became more numerous and increasingly clinically oriented over that span, following a trend that parallels the increasing clinical and translational orientation of both NIH grant applications and NIH-funded publications. Surprisingly, this increase appears to be independent of the representation of clinician-scientists in the NIH workforce.

## Introduction

Biomedical research has significantly contributed to the increased life- and health-spans enjoyed by modern humans. In the last hundred years, physicians and scientists have discovered antibiotics, pioneered the use of insulin as a treatment for diabetes, and developed vaccines against infectious diseases such as polio, HPV, and COVID-19. Immunotherapeutic approaches, including chimeric antigen receptor T cells (CAR-T), have dramatically improved cancer patient survival. The accomplishments of just the past three decades, which include CRISPR-based gene therapy, AI-based protein structure prediction, and the deployment of super-resolution microscopy to describe cellular structures in unprecedented detail, promise further advances. While researchers all over the world contributed to these breakthroughs, the United States has generally been regarded as the global leader in biomedical innovation since the middle of the twentieth century [1–4]. This position has been widely attributed to federal funding support for biomedical science [5], more than 80% of which has been distributed by the National Institutes of Health (NIH; [6]).

A majority of the NIH budget is invested in research project grants (RPGs) held by universities and independent research institutions [7]. RPG funds are distributed through various grant mechanisms (hereafter mechanisms) in support of agency goals. Different mechanisms have different properties [8]. They may be commonly or infrequently used, intended to provide smaller or larger budgets, or designed to support either individual investigators or teams of varying sizes. Some mechanisms allow scientists to submit proposals on a scientific topic of their choice through a parent program announcement (PA); these applications are referred to interchangeably as either unsolicited or investigator-initiated. Other mechanisms are only or primarily used in funding opportunities (historically, funding opportunity announcements, FOA; now Notice of Funding Opportunity, NOFO) published by the different Institutes and Centers of NIH to target specific areas of programmatic priority. Such opportunities can be divided into those with set-aside dollars (Requests for Applications, RFAs, or Program Announcements with Set-aside funds, PAS) and those without (Program Announcements with special receipt, referral, and/or review considerations, PARs); both may be considered an expression of NIH priorities and are therefore considered here as solicitations for applications on a particular topic. A limited number of mechanisms are associated with both unsolicited and solicited funding opportunities.

The original grant mechanism employed by the NIH, and the most widely known, is the R01. A majority of R01 applications are unsolicited, but the mechanism is also used to solicit research proposals on a specific scientific topic defined by the agency. In either case, R01 awards generally support the research of individual scientists. The R01 has also been the subject of most analyses of NIH funding. Previous work has examined the participation of clinician-scientists in R01-funded research [9–13], the increasing age at which scientists obtain their first R01 [14, 15], the challenges that women and underrepresented minorities face in securing an R01 award [16–20], the likelihood of renewing an R01 [21], longitudinal analysis of the productivity of awardees [22], and the increased overall competition for R01 funding [23, 24]. However, there has been relatively little consideration of how R01-funded research fits within the larger landscape of NIH support for biomedical science, or how the position of the R01 within the NIH portfolio may have changed over the years. These questions are particularly salient given that recent decades have seen landmark advances in clinical and scientific knowledge, but the form and purpose of the R01 – to support independent research by individual investigators on a well-defined problem – has remained largely unchanged since at least 1965 [25].

Considering the NIH portfolio as a whole presents a valuable opportunity to analyze whether some administrative structures, because of their size, duration, and/or other unique properties, might be better at delivering biomedical advances. To begin to answer this question, we apply data science methods to analyze the full portfolio of 2,234,898 RPG applications and awards handled by NIH between fiscal year (FY) 1986, the first year for which digitized records are available, and FY2017, the first year in which NIH piloted the broad use of the R35 mechanism to support an entire program of research as opposed to the traditional project-based model exemplified by the R01. We ask how the use of funding opportunities and mechanisms has varied during this period, and whether the types of funding opportunities that provided support for a project relate to the type of publications it produces, which can serve as a proxy for translational progress. We particularly focus on the relationship between the R01 and other mechanisms, and use the National Cancer Institute (NCI), which is the oldest and largest of the twenty-seven Institutes and Centers (ICs) that make up the NIH, to exemplify how long-term funding trends have played out within a single administrative unit. This integrated systems approach allows us to consider NIH funding and the research it supports as parts of a unified whole.

## Results

### An expansion of funding opportunities has driven a surge in non-R01 applications

Before we could determine how different types of grant support contribute to the type of publications produced by awards, we first needed to ask to what extent NIH has used the same mechanisms in the same way over time. The long-standing ubiquity of the R01 might give the impression that the agency’s strategy for disbursing funds has been relatively invariant, but we found this has not been the case. To begin, there has been a considerable increase in the number of funding opportunities offered annually since 1986, the first year for which electronic records are available. Further, this increase has not been uniformly distributed across all grant mechanisms.

Despite the prominence of the R01 mechanism, its use in solicited funding opportunities plateaued in FY1995 (**Figure 1a**). In contrast, and excluding parent or unsolicited announcements, which are a small and relatively constant fraction of the total, the number of funding opportunities offered by NIH over that same time frame using a mechanism other than the R01 has increased over twenty-fold. The total number of non-R01 opportunities peaked in 2007 then dropped by slightly more than a third in 2008, primarily due to a reduction in those using the R21 or R03. This abrupt change was gradually reversed over the subsequent decade through the increasing use of alternate mechanisms, especially the U01.

**Figure 1.**
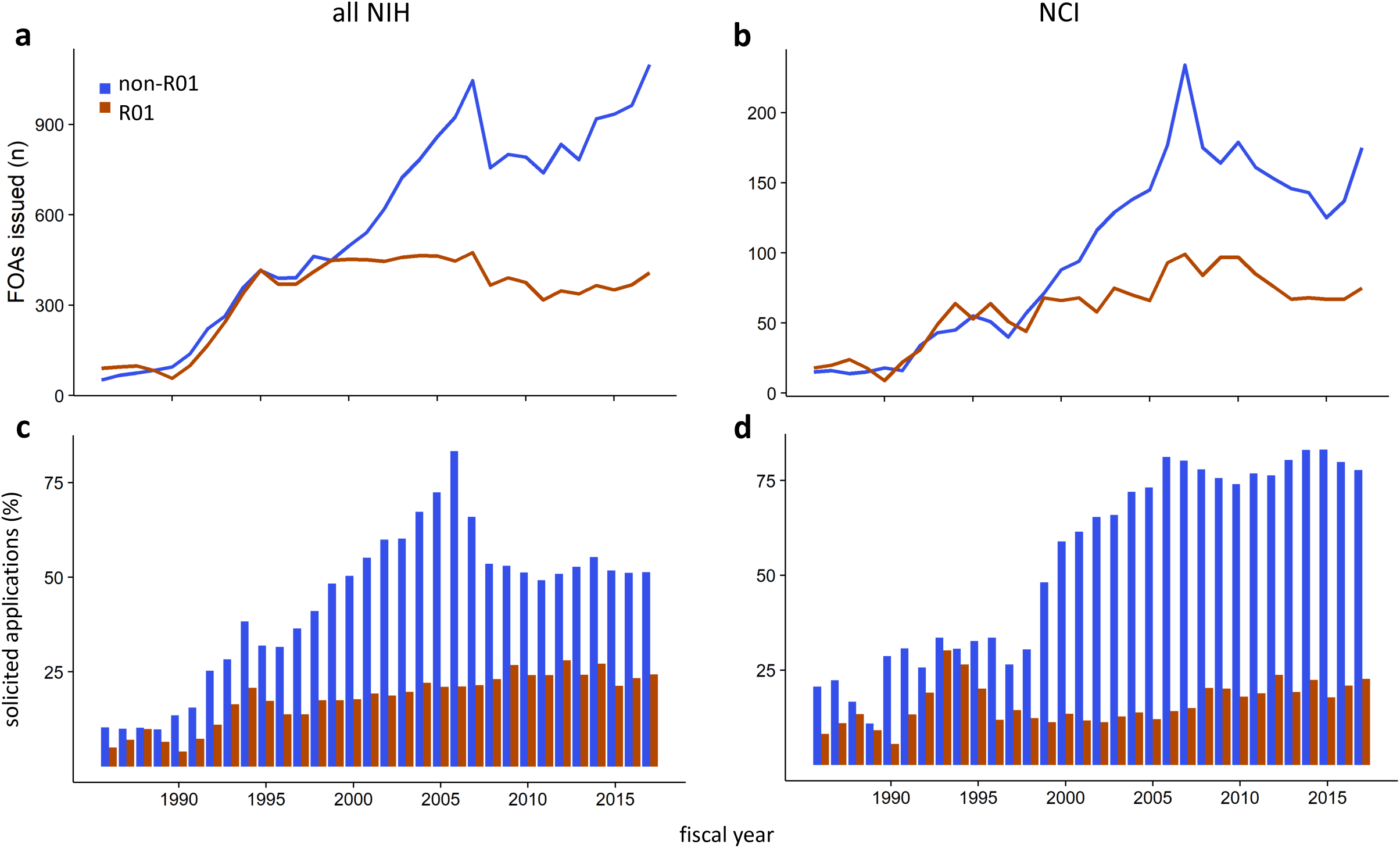
An increase in solicited funding opportunity announcements (FOAs) has driven an increase in applications for non-R01 awards. **(a,b)** Number of solicited FOAs offered each year by NIH (left) or NCI (right) through either an R01 (red) or non-R01 (navy blue) mechanism. **(c, d)** Applications to solicited FOAs as a percentage of all R01 (red) or non-R01 (navy blue) applications received by NIH (left) or NCI (right).

The trends are the same at NCI, although the magnitude and timing of the effects vary slightly; the number of announcements using non-R01 mechanisms increased eight-fold and overtook the number of R01-based programs in FY2000 (**Figure 1b**). Interestingly, R03 and R21 opportunities gradually returned at NCI while the number of U01 offerings also grew.

Both across NIH and at NCI, the fraction of non-R01 applications received in response to a solicitation increased largely in parallel with the number of open funding opportunities (**Figure 1c-d**). Importantly, this response is not driven by increased interest in programs with dedicated or set-aside funding, since these constitute no more than a third of solicited announcements (**Supplemental Figure 1a-b**), and the fraction of applications received under them has been small and stable over more than thirty years (**Supplemental Figure 1c-d**).

Given that inflation-adjusted budgets have remained constant or increased only very slightly in size (**Supplemental Figure 2a-d; Supplemental Data 1**), and that award rates for R01 and non-R01 mechanisms have been roughly equivalent since the early 2000s (**Supplemental Figure 2e-f**), the raw increase in number of non-R01 applications (**Figure 2**) is sufficient to explain the modest shift in the NIH portfolio away from R01 and towards non-R01 awards (**Supplemental Figure 3**). This previously unappreciated alteration in the way research is supported highlights the importance of understanding both how these relatively under-analyzed mechanisms are used, both by the agency and applicants, and their net effect on scientific outputs.

**Figure 2.**
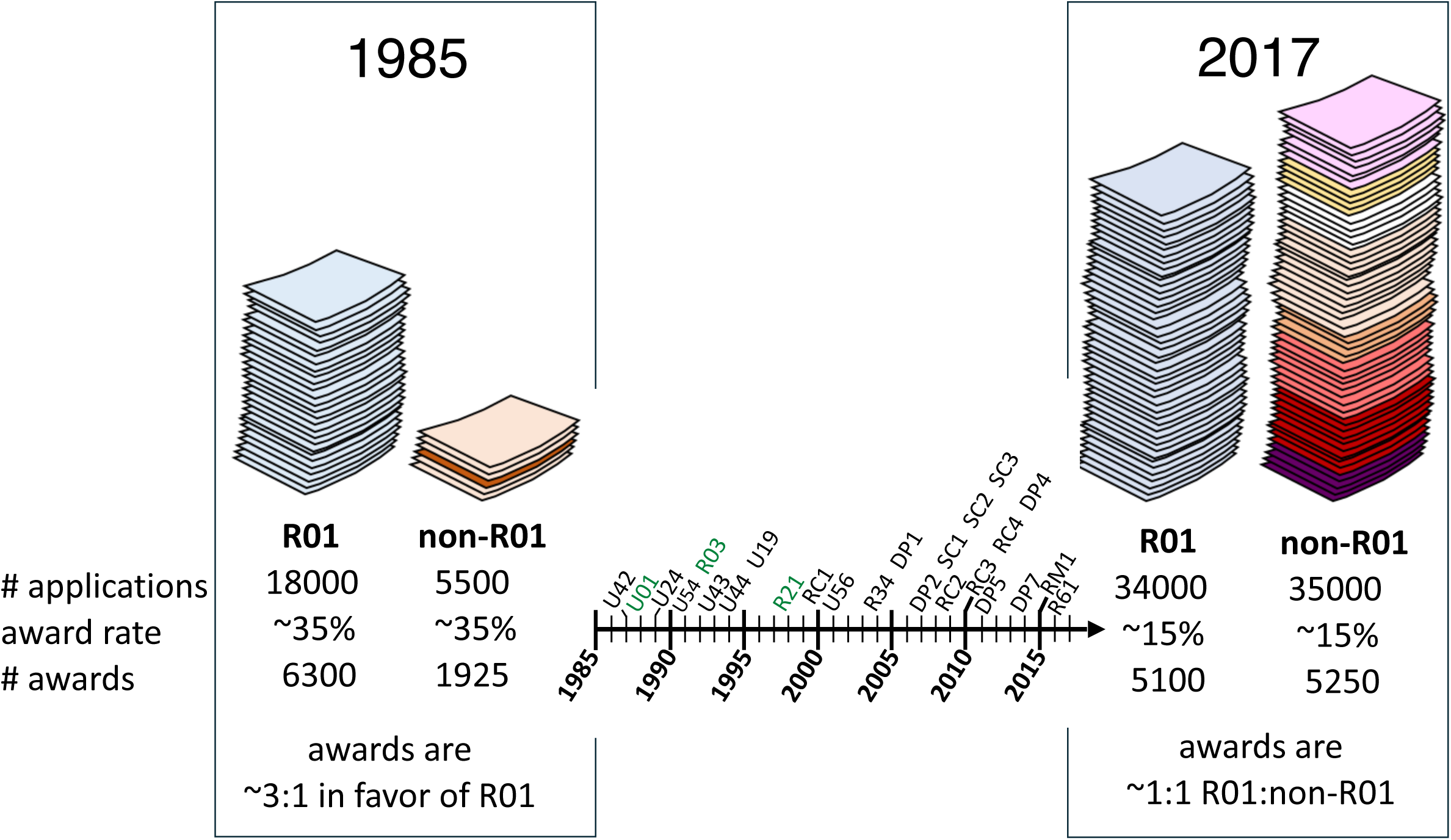
Proliferation and expansion of non-R01 mechanisms is sufficient to explain the shifting composition of the NIH RPG portfolio. Left side, similar award rates combined with a differential number of applications for R01 (light blue) and non-R01 (orange) awards result in a portfolio that favors R01s by approximately 3 to 1. Right side, similar award rates and similar application numbers result in a portfolio in which R01 (light blue) and non-R01 (multicolored) awards are equally prevalent. Center, timeline showing first documented use of a new mechanism (black text) or, for mechanisms that predate the timeframe of this analysis, the first time NIH received more than 1000 applications per year through that mechanism (green text).

### A majority of the growth in the portfolio can be attributed to a subset of non-R01 mechanisms

Excluding the R01, our dataset contains a total of 96 unique mechanisms (**Materials and Methods**). However, not all of these are equally well represented. The most commonly occurring single mechanism, both in terms of opportunities offered and applications received, is the R21. A substantial fraction of non-R01 announcements – in the peak year, close to 30% across NIH and exceeding 50% at NCI – made use of the R21 mechanism (**Supplemental Figure 4**). R03 and U01-based announcements also become increasingly common over the time frame of our analysis. Together, R21, R03, U01, and R01 mechanisms account for 60 to 75% of announcements offered by NIH, and 60 to 95% by NCI, depending on the year in question (**Supplemental Figure 4**).

In terms of applicant response to funding opportunities published across NIH, the R21 again dominates, with the number of applications received increasing by more than 150-fold since the early 1990s (**Figure 3a, Supplemental Table 1**). Applications for R03 and U01 awards each increased approximately 4-fold, consistent with the frequent use of these mechanisms in funding opportunities; applications for R15 awards also increased by more than 5-fold (**Figure 3b-d**), despite the number of funding opportunities for that mechanism remaining consistently small. However, not all mechanisms saw a large uptick in interest over time. The number of R01 applications increased less than two-fold, while applications for P01 grants, a mechanism intended to support knowledge and resource sharing among a group of independent investigators working on related projects, have remained stable or trended slightly down (**Figure 3, e-f**). Similar trends occurred at NCI (**Figure 3, g-I; Supplemental Table 2**).

**Figure 3.**
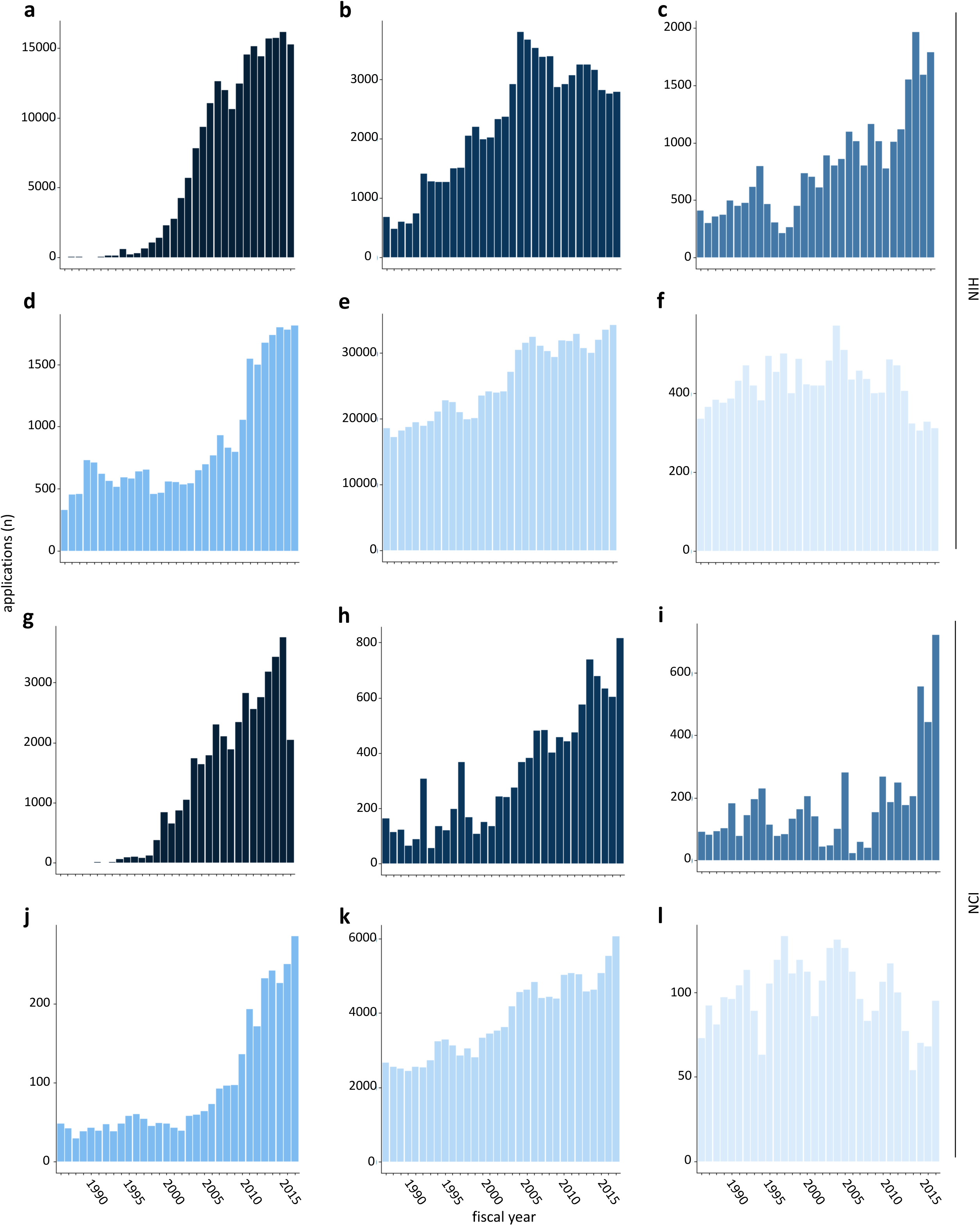
Increase in the number of non-R01 applications. Number of applications for the most common non-R01 research grant mechanisms received by the NIH (top six panels) and the NCI (bottom six panels) between fiscal years 1986 and 2017. (**a, g**) R21 (**b, h**) R03 (**c, i**) U01 (**d, j**) R15 (**e, k**) R01 (**f, l**) P01.

In this context, the growth of the U54 mechanism, which supports multidisciplinary and often multi-institutional research centers through large awards, stands out as a particularly interesting case. Like the P01, the U54 supports resource sharing and collaboration. Unlike P01s, applications using this mechanism have grown approximately 60-fold NIH-wide since its introduction in 1991 (**Supplemental Figure 5a; Supplemental Table 1**). Although the U54 can in theory be used to support any part of the full range of research and development from the most fundamental to clinical, we noted that in practice, funding opportunities that use it are focused on clinical and translational science, as indicated by the frequency with which those keywords appear in their titles. The related keywords human, disease, health, health disparities, and cancer are also common (**Supplemental Figure 5b**). Consistent with the frequent appearance of the keyword “cancer”, the NCI has offered more funding opportunities and received more competing applications through this mechanism than any other individual Institute or Center of NIH (**Supplemental Figure 5c, d**).

### Non-R01 mechanisms have been a driver of the increase in clinical and translational research

To determine to what extent other non-R01 mechanisms focused on clinical and translational research, we first applied the Medical Subject Headings (MeSH) vocabulary developed and maintained by the National Library of Medicine (NLM) to the text of funding announcements offered between 1992, when the NIH Guide to Grants and Contracts was first digitized, and 2017, the end of the time frame analyzed here. All MeSH descriptors can be assigned to one of three broad areas: Molecular and Cellular, Animal, or Human. Previous work has defined an increase in Human-related MeSH terms as a marker of translation [26, 27]. Controlling for the overall length of the text and the number of announcements offered annually, there has been a significant increase in the use of Human terms in non-R01 based announcements; in contrast, R01-based funding announcements have changed very little (**Figure 4a, b**). Terms in the Animal category have remained uniformly rare, while those in the Molecular and Cellular category have become slightly more common in non-R01 based announcements (**Figure 4c-f**).

**Figure 4.**
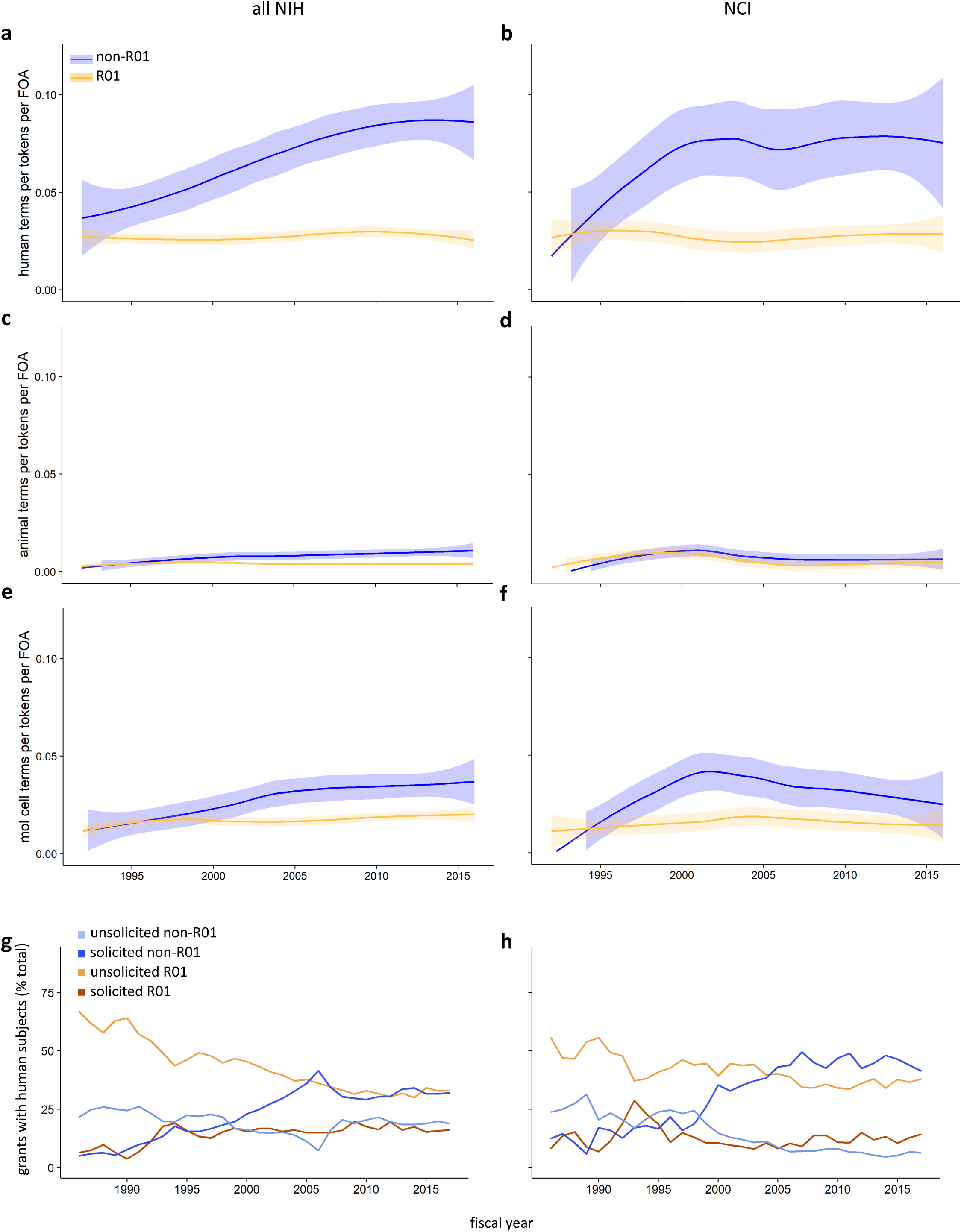
Increasing clinical and translational orientation of non-R01 funding opportunities and applications. (**a-f**) Increasingly human-centered language in the text of NIH funding opportunity announcements (FOAs). Number of terms from the human (**a, b**), animal (**c, d**), or molecular and cellular (**e, f**) branch of the MeSH tree, normalized to the number of terms per FOA and the number of FOAs per year, for NIH (left) or NCI (right). R01, orange; non-R01, blue; shaded area represents the 95% confidence interval. (**g, h**) Shift in the type of grant applications that propose work with human subjects. Unsolicited R01 (orange), solicited R01 (dark orange), unsolicited non-R01 (light blue), and solicited non-R01 (dark blue) applications with human subjects as a percentage of all (**g**) NIH or (**h**) NCI applications with human subjects.

Consistent with this change, proposals that include human subjects, another proxy for research with a clinical and translational focus, have shifted from predominantly unsolicited R01 applications in the 1990s to a close to even mix of unsolicited R01 and solicited non-R01 submissions in recent years, both across NIH and at NCI, over the same time frame (**Figure 4g, h**).

### NIH-funded publications show an increased clinical and translational orientation

The changing number and types of programs designed to support clinical and translational science led us to ask whether there was any downstream change in the research papers produced by these grants. To answer this question, we analyzed ∼2.3M papers published between 1986 and 2017 that cite NIH grant support. We calculated a Human MeSH score, which is a function of the number of terms in the human branch of the MeSH ontology, for each individual paper. We also determined what fraction of the 2.3M papers were flagged each year by PubMed as clinical articles (see **Materials and Methods** for details). Our data show that NIH- and NCI-supported work has become more clinically oriented overall, as measured by either average human MeSH score or publication of clinical articles (**Figure 5a-d**). Fractionation of these two metrics across all grants acknowledged by an individual paper shows this trend is being driven almost entirely by support from solicited non-R01s. Across NIH, unsolicited awards have provided decreasing support for clinically oriented papers, regardless of mechanism (**Figure 5e, g**). At NCI, clinical articles have consistently acknowledged only low levels of support from any type of R01, but the average human MeSH score of papers attached to unsolicited R01s has dropped sharply since 2008 (**Figure 5f, h**).

**Figure 5.**
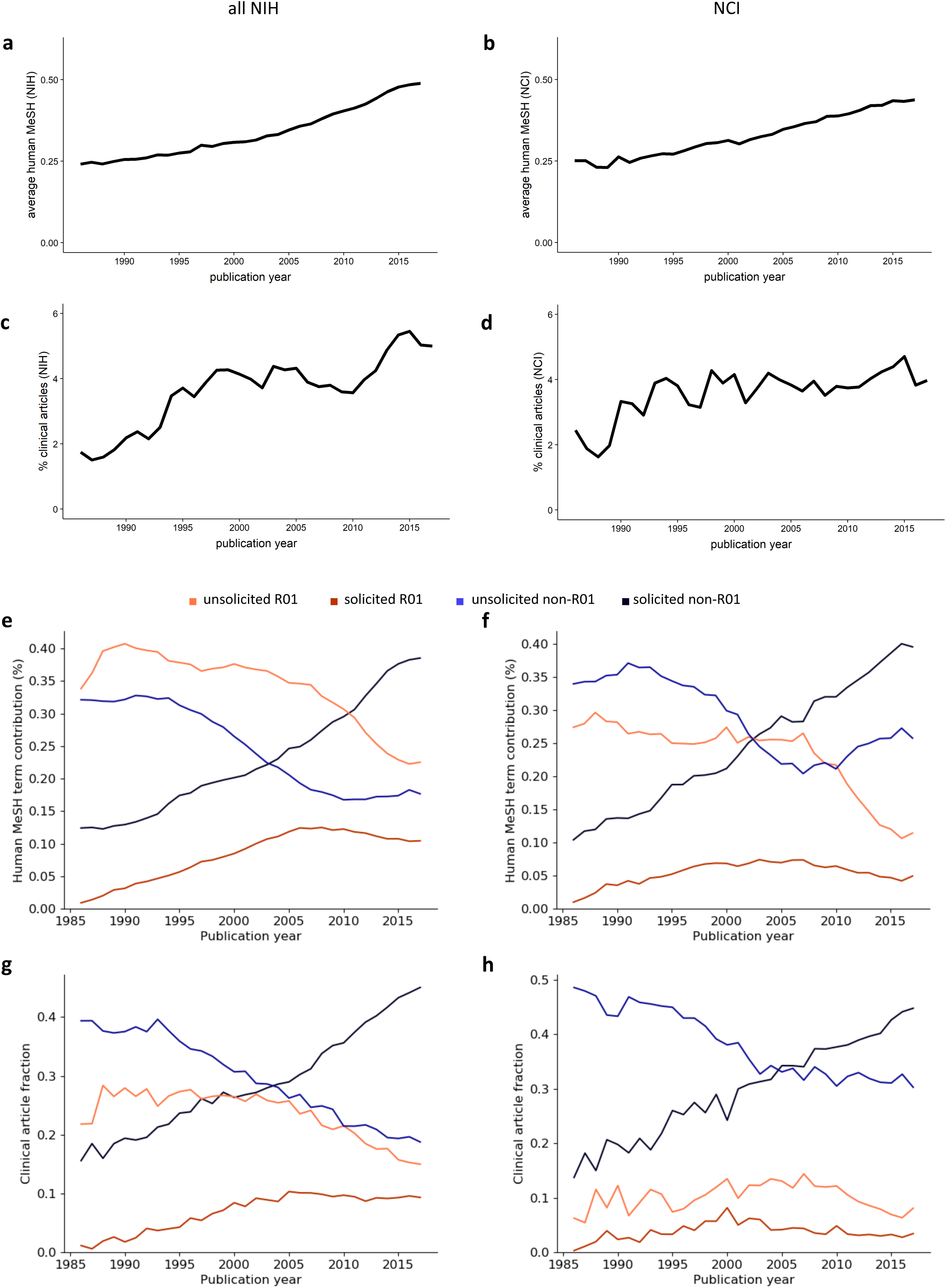
Increasingly clinical and translational orientation of NIH-funded publications. (**a, b**) Increase in the average human MeSH score of papers that cite support from NIH or NCI awards. (**c, d**) Increase in clinical articles as a fraction of all papers that cite support from NIH or NCI awards. (**e, f**) Fraction of human MeSH score supported by unsolicited R01 (orange), solicited R01 (red), unsolicited non-R01 (blue), and solicited non-R01 (navy) awards made by NIH or NCI awards. (**g, h**) Fraction of clinical articles supported by unsolicited R01 (orange), solicited R01 (red), unsolicited non-R01 (blue), and solicited non-R01 (navy) awards made by NIH or NCI awards.

We next considered the possibility that the increase in both clinically oriented papers and human-focused terminology in funding opportunities might be coincidental, or a product of larger shifts in the research landscape beyond and outside the influence of NIH. To begin to address this question, we measured the clinical orientation of more than 610,000 PubMed-indexed papers, approximately half of which cite NIH support. We found that both the average human MeSH score and the Approximate Potential to Translate (APT), the latter of which predicts the likelihood of citation by a clinical article [27], have also increased specifically for NIH-supported papers (**Figure 6a, b**). The fraction of papers flagged by PubMed as clinical articles shows a more complex pattern. This article type increased in both groups through 1999 but diverged thereafter, becoming more common among papers acknowledging NIH support but less common among those that do not, so that the two groups are indistinguishable by the end of the time period in question (**Figure 6c**). On the whole, however, and consistent with another recent report [28], these three measures indicate that the end products of NIH awards have been increasing in their clinical and translational orientation, and suggest a gradual but wholesale shift in the focus of the agency.

**Figure 6.**
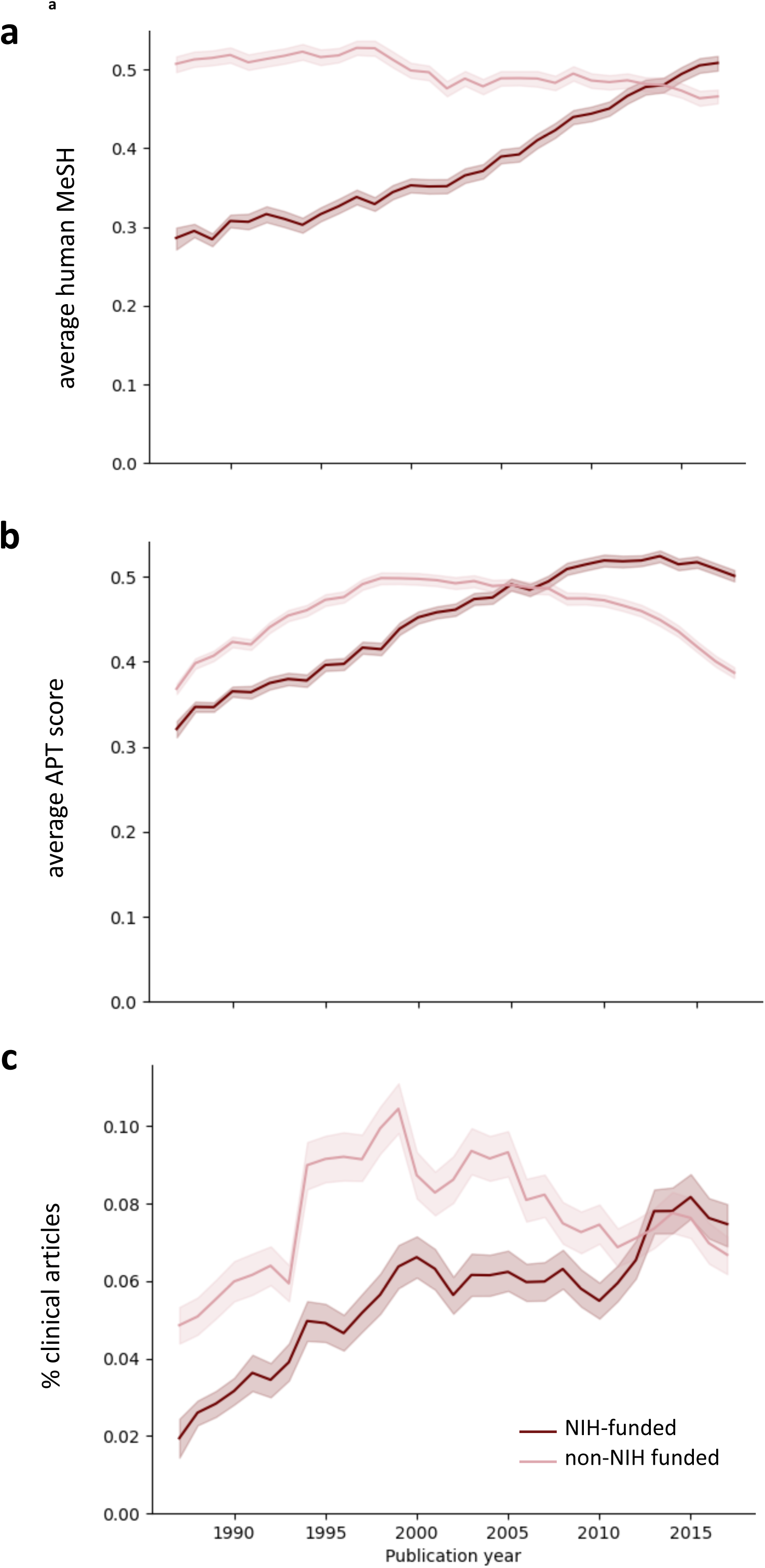
Comparison of the clinical orientation of NIH-funded and non-NIH funded publications. (**a**) Average human MeSH score; (**b**) APT scores; and (**c**) clinical articles as a fraction of all papers that cite support from NIH awards (red) or another non-NIH funding source (pink).

### Participation of clinician scientists in the NIH workforce

Previous studies have acknowledged that many NIH-supported investigators who pursue clinical research do not hold a medical degree [29, 30]. At the same time, there has been a longstanding recognition that physician-scientists hold a special position in the conduct of clinical research, primarily due to their status as caregivers for human subjects participating in interventional studies [31, 32]. As a result, concerns about the size and success of the physician-scientist work force have frequently been considered alongside concerns about support for clinical research [10, 29–36]. Two observations—that the pipeline of NIH research, from upstream funding opportunities to downstream award products, has turned increasingly towards clinical and translational science, and that practitioners of at least one medical specialty, surgery, have increased their representation in the NIH workforce over the latter third of the time frame considered here [9]—led us to wonder whether overall physician participation in the NIH workforce also increased. Strikingly, we found that since 1992, the percentage of investigators with an MD has remained within a standard deviation of the mean value calculated over the time frame of our analysis (**Figure 7a**). Approximately half of the modest growth in the population of clinician-scientists comes from investigators with both an MD and a PhD; while the absolute number of these dual-degree holders remains small, their relative representation has grown slightly more than three-fold. In contrast, the percentage of investigators who are solely MD holders peaked in 2006 and declined continuously from 2010 onwards (**Figure 7b**). These data indicate the factors influencing the production of more clinically focused research outputs are complex, and do not necessarily include a requirement for further growth in the total pool of clinical investigators.

**Figure 7.**
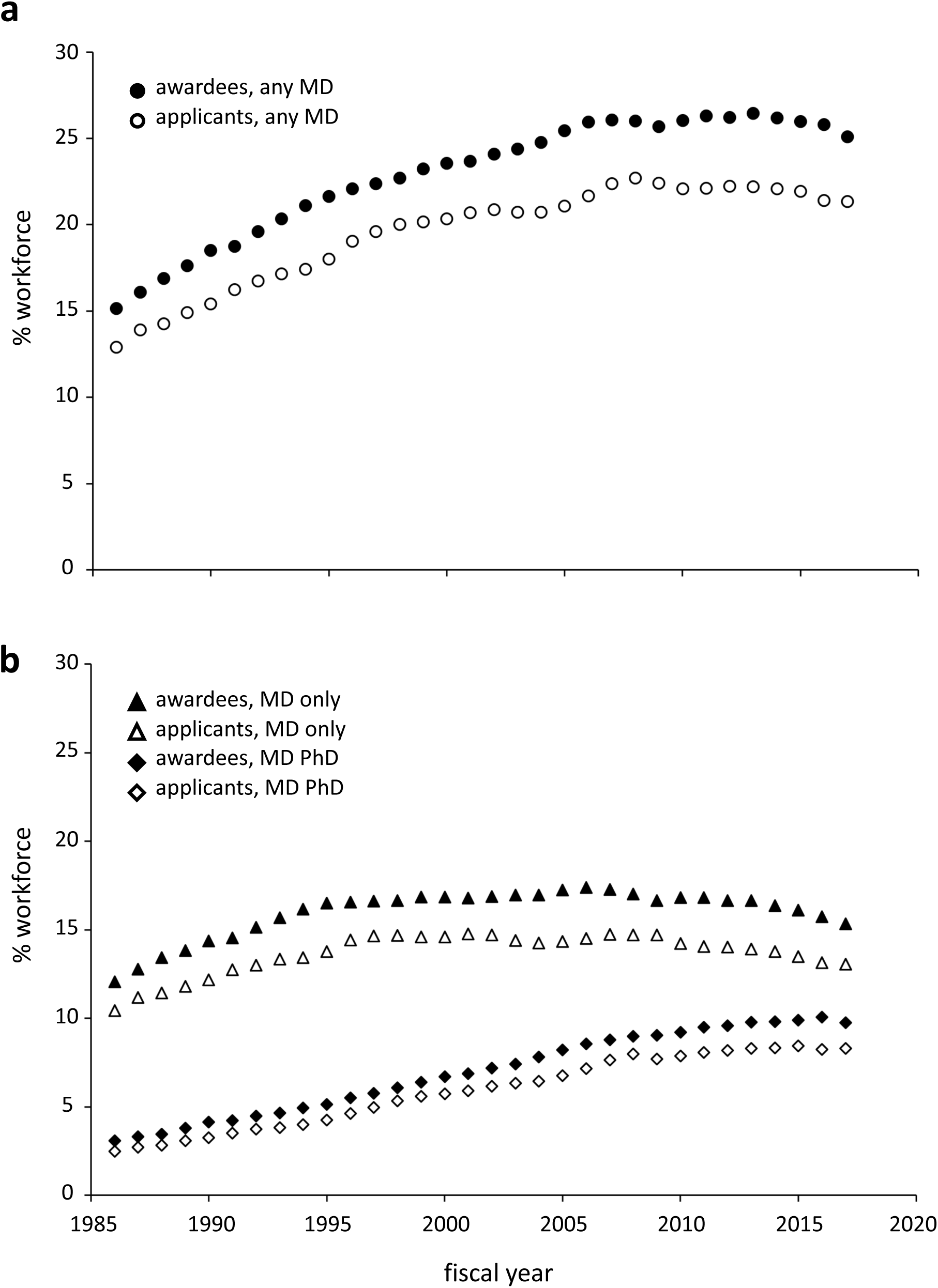
Representation of physician-scientists in the NIH workforce. (**a**) Percent of all awardees (filled circle) or all applicants (open circle) that listed their degrees as including an MD when submitting an NIH proposal. (**b**) Percent of all awardees (filled symbols) or all applicants (open symbols) that listed only an MD (triangle) or both an MD and PhD (diamond) when submitting an NIH proposal.

## Discussion

We show here that research supported by NIH moved in a more clinical direction over a span of 32 years, in parallel with a shift to more clinically oriented funding announcements and a reduction in R01s as a fraction of all RPGs. Three lines of evidence suggest these trends may be connected. First, the text of non-R01, but not R01, based funding opportunities has become increasingly human focused over time (**Figure 4a-f**). Second, studies that include human subjects have largely shifted from unsolicited R01s to solicited non-R01s (**Figure 4g, h**). Finally, solicited non-R01s have become the primary source of support for clinically and translationally oriented publications (**Figure 5e-h**). These changes cannot be attributed solely to expansion or tightening of the NIH budget, a phenomenon referred to by economists as budgetary shock, since they began prior to and continued past the end of the 1998-2003 budget doubling and did not appear to be affected by sequestration in 2013.

It is difficult to determine exactly what brought about the parallel shifts reported here, and multiple factors may be at play. NIH-supported publications specifically were becoming progressively more clinically and translationally oriented as far back as the late 1980s. Since the text of FOAs is only available electronically from 1995, our method cannot definitively establish causality from the chronology of events. The clearest discontinuity in what appears to be overall gradual trends occurred in the year 2000, when the number of solicited funding opportunities offered using non-R01 mechanisms diverged decisively from those using the R01 (**Figure 1a**). At least three significant events occurred around this time, any or all of which could have played a role: the publication of the first draft sequence of the human genome in 1999 [37]; the planning and implementation of a reorganization of the Center for Scientific Review in 2000, two explicit goals of which were to facilitate translational progress [38] and to increase the representation of clinicians on panels [30]; and the public announcement in 2003 of the NIH Roadmap for Medical Research, which aimed to completely re-organize and expand clinical research in the United States [39]. Regardless of the exact nature of the drivers, future investigations of the impact of NIH funding over an extended time frame must consider the shifts described here, namely the movement of NIH-funded research in a more clinical direction and the parallel reduction in R01s as a percentage of RPGs.

Since 1945, the federal research enterprise in the US has operated on the principle that freedom of inquiry and the pursuit of intellectual curiosity are essential to scientific progress [40]. This foundational doctrine could be undermined if applicants feel pressure to redirect their research in a more clinical direction. Moreover, proposals that are conceived as investigations into fundamental biology are unlikely to be strengthened by the addition of tangential clinical or translational experiments. At the same time, the continuity and stability of federal research funding depend (appropriately) on demonstrated benefit to the public [40]. No individual scientist can balance this equation on their own; an organized effort is needed to set priorities and ensure those priorities are effectively implemented through available funding opportunities.

We chose to end our analysis in 2017, the year R35 Outstanding Investigator Awards designed to support an investigator’s entire research program, rather than an individual project, were first widely offered to applicants to the National Institute of General Medical Sciences (NIGMS) as a replacement for the R01. Although other Institutes and Centers, most notably NCI and the National Heart, Lung, and Blood Institute (NHLBI) also later offered this option to a subset of their applicants, since 2017 a supermajority of R35 awards have been granted by (NIGMS) and therefore fall into the category of fundamental research. This new funding paradigm represents a departure from NIH’s historical approach, and its inclusion here could confound the identification and/or interpretation of longer-term trends in the support for and production of research to improve human health. Future targeted analysis will be needed to determine both the effectiveness of the R35 mechanism and its impact on the direction of the NIH portfolio.

It is important to note that while our work identifies clear overall trends in the NIH and NCI portfolios, individual funding opportunities are highly variable in their scientific scope, goals, and effectiveness. Clinical and translational research is also a heterogenous category. As we have defined it here, it includes epidemiology, behavioral science, clinical practice, diagnostics, precision medicine, and more. While it is both informative and important to consider this very high-level perspective, it should not be assumed that our data accurately represents the situation at more granular levels. Each scientific field in biomedicine changes size at its own rate, encounters its own challenges, and makes progress in its own time. Understanding the state of any individual field requires dedicated analysis. Similarly, determining whether a specific funding opportunity achieved its goals requires dedicated analysis of that funding opportunity. Our data should not be taken as an indication that all funding opportunities are equally successful, nor should it be used in the absence of other evidence to support the assumption that an individual funding opportunity must have had some scientific impact.

As is true of research areas and funding opportunities, there is significant heterogeneity within the larger category of non-R01 mechanisms. Individual mechanisms may or may not follow the same pattern of growth we see for the category as a whole and will vary in their orientation towards clinical topics. The R21, which has experienced the greatest increase in both funding opportunities and applications (**Supplemental Figure 4;** **Figure 3a** and **g**), illustrates this complexity. Although the R21 supports research across the entire fundamental to clinical spectrum, previous analysis has shown that R21 awards to surgeon scientists were more likely to be clinically oriented in 2020 than in 2010 [9], suggesting the mechanism may be playing a role in both trends. While further work is needed to characterize the scientific topics supported by the R21 and to determine how these topics may have changed over time, an in-depth analysis of this single, specific mechanism is beyond the scope of this work.

The number of funding opportunities offered using a particular mechanism obviously factors into, but is insufficient to explain, the number of applications generated in response. For example, while the 150-fold NIH-wide increase in R21 applications corresponds to a roughly 100-fold increase in unique R21 funding opportunities, a 30-fold increase in R03 opportunities correlates to a much more modest 4-fold increase in proposals. Since both are two-year non-renewable awards, the roughly 3-fold difference in award size ($275,000 versus $100,000 maximum) is a likely driver in their differential effect. However, award size alone is also insufficient to explain applicant response. Applications for P01s, large awards that support multiple investigators and dedicated core facilities, remained remarkably constant in an environment where application numbers increased overall. Given the widespread sentiment that biomedical research is becoming more collaborative and interdisciplinary [41, 42], it is somewhat surprising that there has been no real growth in the original mechanism intended to support broadly based, synergistic, and multidisciplinary work. This may be an example of applicants responding to NIH actions, as relatively few P01-based funding opportunities have been offered by the agency. Alternatively, it may be because some aspect of the P01 application process is inhibitory, because a relatively small portion of the workforce is in a position to respond to opportunities that require synergistic interactions between a multi-investigator team, and/or because applicants believe they may improve their chance of award by targeting a U54 or other U-based funding opportunity designed to support collaborative research in an area targeted by NIH.

One surprising aspect of the data presented here is the demonstration that applicants respond strongly to the announcement of opportunities that are not accompanied by set-aside funding. During this time period, approximately two-thirds of all applications were received in response to solicited funding opportunities with no set-aside funds. This observation runs counter to the assumption that the availability of such funding provides a strong incentive that will effectively increase submissions. It is consistent, however, with previous work demonstrating that where set-aside funds are available, applicants generally constrain themselves to programs that are a close match to their scientific expertise; the likelihood an investigator will submit an application decreases as the scientific topic of that program becomes more distant from their previously published work [43]. This selective behavior on the part of applicants, taken together with the fact that funding opportunities attached to set-aside funds are rare and tend to be offered for shorter durations, means that, on the whole, programs without set-aside funds actually have a much greater impact on the overall direction of funded research.

Another counter-intuitive observation we report is an increase in the production of clinically-oriented literature without an accompanying increase in the representation of clinician-scientists in the NIH workforce, either as applicants or as awardees. There are a number of possible explanations for this, none of which are mutually exclusive. The average number of papers produced per funded clinician may simply have increased. There may have been an increase in collaboration, with MDs participating in roles other than that of lead investigator. It is also possible that the increasing footprint of MD-PhD scientists in the investigator pool has had a meaningful impact on the direction of research outputs. Finally, it may be that efforts to increase the representation of clinicians on review panels [30] have influenced the direction of funded research, and subsequently the direction of research outputs, for all investigators. None of these hypotheses lessen the importance of the role clinicians play in caring for patients participating in trials, but they do highlight the need for a better understanding of how the clinical perspective contributes to research-driven improvements in human health.

Future analyses are also needed to determine whether funding opportunities have been effective in driving fundamental scientific progress, which is related but not identical to translational progress. Since its earliest days, NIH has recognized that understanding the elemental principles of biology is intertwined with making improvements to human health and lifespan [44, 45]. However, while it is generally agreed that both fundamental and clinical research are necessary to produce medical advancements, the optimal balance between these two types of investigations has been debated for decades, and remains unknown. Large-scale study of the trajectories of past biomedical successes may help answer this question going forward. Regardless of the answer, overall trends in support for different scientific fields should be monitored, and particular care should be taken to ensure that any changes of direction are fully intentional.

Setting an overarching direction for future research that further improves human health requires an understanding of the path between the current state of knowledge and the desired goal. As in other areas of human endeavor, the past is often a helpful guide to the future. A data-driven understanding of cause and effect can support our understanding of what factors led to past advances. This may, in turn, help funders and scientists identify future advances at an earlier stage and take steps to speed their progress. Retrospective analyses also promote transparency and are key to determining how well the intention of a program matched its actual, real-world effect. Additional analyses of the portfolios of both public and private funders of biomedical science could yield invaluable insights, fostering a more informed and strategic approach to funding that enhances the efficiency and effectiveness of research aimed at improving human health.

## Methods

### Data sources

Data on 2,234,898 (including 1,443,458 competing, i.e. Type 1 and Type 2) grant applications and awards submitted to NIH between 1986 and 2017 were extracted from the Information for Management, Planning, Analysis, and Coordination (IMPAC II) database, which is used by NIH staff to track and manage research grants and contracts. To provide maximal context, the category of non-R01 mechanisms was defined broadly, to include the D43, DP1, DP2, DP3, DP5, DP7, G08, G12, M01, OT2, OT3, P01, P20, P2C, P30, P40, P41, P42, P50, P51, P60, PL1, PN1, PN2, R03, R15, R18, R21, R22, R23, R24, R25, R28, R29, R33, R34, R35, R36, R37, R41, R42, R43, R44, R50, R55, R56, R61, R90, RC1, RC2, RC3, RC4, RF1, RL1, RL2, RL5, RL9, RM1, S06, S07, S10, S11, S21, S22, SB1, SC1, SC2, SC3, U01, U10, U18, U19, U24, U2C, U2R, U34, U41, U42, U43, U44, U54, U56, UA5, UC1, UC4, UC7, UF1, UG1, UG3, UG4, UH1, UH2, UH3, UL1, UM1, and UM2 activity codes. To avoid double counting, subprojects were excluded. ARRA applications and awards were also excluded. Applications were identified as unsolicited based on the presence of an IMPAC II flag indicating they were either unsolicited (prior to FY2007) or were received under the parent funding opportunity announcement (since FY2007); all other applications were considered as solicited. Data on awards are publicly available through the NIH RePORTER (https://reporter.nih.gov/). Per NIH policy, data on unawarded applications are considered confidential and protected from disclosure other than for purposes of evaluation. Data, including metadata, html source files, title and text, for active and expired Funding Opportunity Announcements (FOAs) published between 1992 (the earliest year for which digitized records are available) and 2017 were downloaded from the NIH Guide for Grants and Contracts (https://grants.nih.gov/funding/searchguide/index.html#/). Data and metadata on publications and their funding sources were extracted from PubMed (https://pubmed.ncbi.nlm.nih.gov/). Supplemental data for this work are available at https://figshare.com/.

### Analysis of award budgets

To quantify trends in award size over time, the Biomedical Research and Development Price Index (BRDPI)-adjusted average total costs of competing (T1 and T2) grants in each year of our analysis was subjected to a Mann-Kendall test using the pymannkendall 1.4.3 package. Awards with a budget of <=$1000 were dropped, as these records very likely reflect post-award actions such as transfers or terminations that were taken by grants administration. Trends were calculated for NIH as a whole, and separately for each of six large Institutes, for non-R01 and R01 awards. R01s were then further broken down into solicited and unsolicited categories.

### Content analysis of Funding Opportunity Announcements

The set of RPG-based NIH FOAs analyzed here was defined by filtering out any items in the NIH Guide for Grants and Contracts that were assigned to a training mechanism (all T, F, and K mechanisms) or that were issued by an administrative unit outside of NIH, such as the Food and Drug Administration (FDA) or Centers for Disease Control (CDC). The Purpose, Program, Research Objectives, and Background sections were parsed and extracted from FOAs offered in years 1992-2002. For years 2002-2021, the Table of Contents was parsed and Section I, which contains the Background and Research Objectives subsections, was extracted. Extracted content was then tokenized and MeSH terms were isolated using a named entity recognition algorithm. To control for potential differences in the length of the text, the frequency of Animal, Human, and Molecular/Cellular terms was then normalized to the total number of tokens extracted from that particular FOA, then the number of terms per token was normalized to the total number of either R01 or non-R01 Funding Opportunity Announcements offered per year. Code is available at https://github.com/NIHOPA/foa.

### Analysis of the degree of clinical orientation of the biomedical literature

Publications are classified as Human, Animal, Molecular/Cellular Biology, or a combination of these three, according to Weber’s algorithm for using the MeSH terms [26], which are assigned to publications by NLM indexers [46]. MeSH terms persist once assigned, although additional terms may be added over time as the MeSH vocabulary expands. The degree to which a paper is clinically oriented was measured based on the fraction of its associated MeSH terms that are assigned to the human branch of the MeSH ontology. For each paper, this human MeSH score was then divided equally across all grants (including training grants) cited as support. The human MeSH score was summed for all papers in each publication year in the dataset, and the fraction attributed to grants in each of four categories (solicited R01, unsolicited R01, solicited non-R01, and unsolicited non-R01) was plotted relative to the total human MeSH score for all supported papers in that year. As previously described [27], a paper was designated a clinical article if PubMed identified it as a clinical trial, clinical study, or clinical guideline.

### Comparison of NIH and non-NIH funded biomedical literature

A random sample of NIH-funded papers (n = 302,138) was compared to non-NIH funded papers (n = 308,086). To avoid any indirect effects of NIH policy on the research directions of US academics who might be pursuing funding from the agency, non-NIH funded papers were randomly selected from primary research articles written in the English language with a Relative Citation Ratio (RCR) of at least 1.0 whose location fields include Europe, Asia, or Australia but NOT United States or North America. For this analysis, papers attributed to NIH were selected from the subset of supported works that can be identified by a fuzzy string match to an NIH award self-reported by the recipient in the Acknowledgements section of the paper [47].

### Identification of clinician scientists in the applicant and awardee pools

Degree information self-reported by contact PIs was used to identify unique clinician applicants and awardees in our dataset. Clinician-scientists were those for whom the degs.deg_code field in IMPAC II included a match to the string “MD”, either alone or in addition to a PhD or other degree; DMD (dentistry), VMD (vet), and PHMD (pharmacist) degrees were explicitly excluded from the string match. For the purposes of this analysis, all contact PIs not identified as MDs or MD, PhDs were designated as non-clinician principal investigators. For each year, unique PIs were then split into four groups: those MD-awarded, MD-not_awarded, not_MD-awarded, and not_MD-not_awarded. These categories were mutually exclusive: a PI with at least one award in that year was counted in the awarded category, and one reporting an MD degree at least once was counted in the MD category. We then plotted the fraction of NIH RPG applicants and awardees who were MDs. If an applicant received an award in a given year, they are counted as an awardee, regardless of whether they submitted one or more than one application.

## Acknowledgements

We thank Chris Pickett of the Office of Portfolio Analysis and members of the Division of Cancer Biology at NCI for their thoughtful feedback. Funding: The authors received no specific funding for this work, but all were employees or contractors for the NIH. Author contributions: Conceived and designed the analysis: K.A.W. and G.M.S. Project management/oversight: K.A.W. and G.M.S (with special thanks to Rebecca Meseroll). Data management and acquisition: B.L.B., J. M.T., and K.A.W. Analyzed the data: B.L.B., J.M.T., and K.A.W. Contributed to writing/editing of the paper: B.L.B., S.E.A., K.A.W., and G.M.S.

## Notes

https://github.com/NIHOPA/foa

https://figshare.com/projects/A_thirty_year_trend_of_increasing_clinical_orientation_at_the_National_Institutes_of_Health/268799

### Competing Interest Statement

The authors have declared no competing interest.

