## Supplemental Figures and Tables for "A thirty-year trend of increasing clinical orientation at the National Institutes of Health"

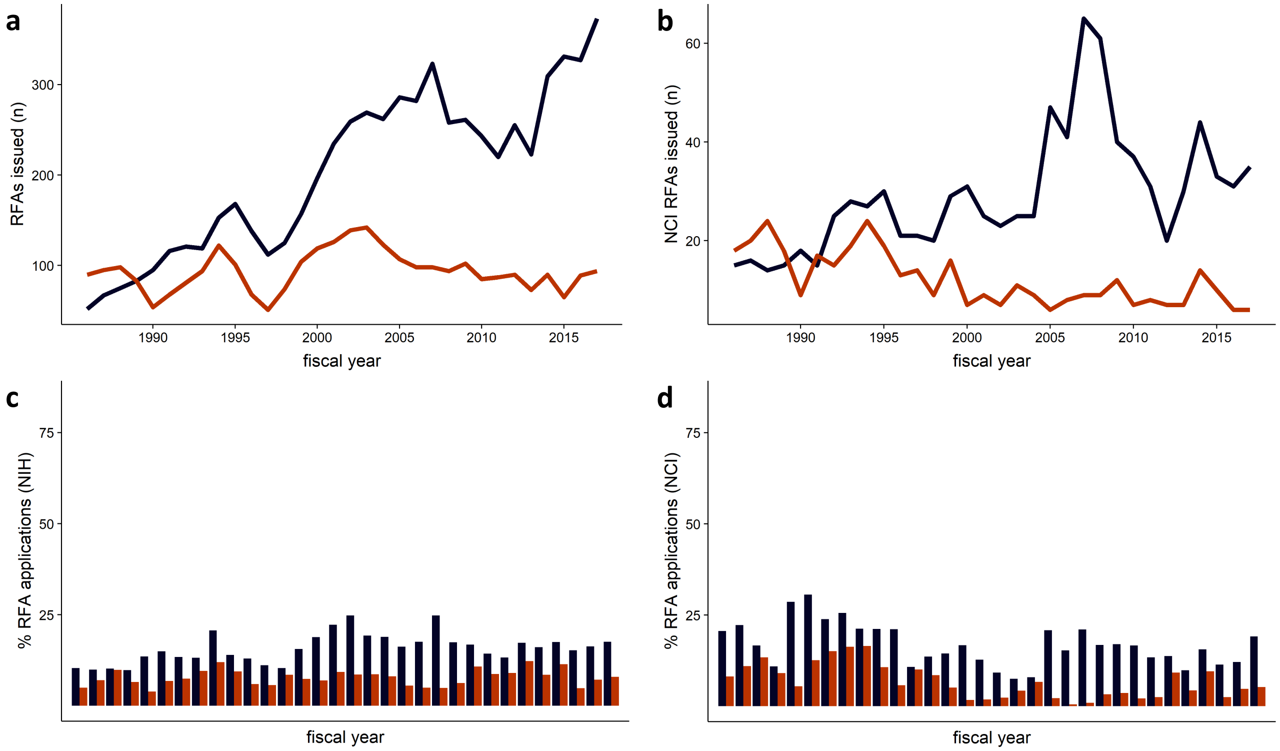

**Figure S1. Minimal effect of funding opportunity announcements (FOAs) with set-aside dollars on receipt of applications.** (**a, b**) Number of FOAs with set-aside dollars issued each year by NIH (left) or NCI (right) through either an R01 (red) or non-R01 (navy blue) mechanism. (**c, d**) Applications to FOAs with set-aside dollars as a percentage of all R01 (red) or non-R01 (navy blue) applications received by NIH (left) or NCI (right).

Figure S1

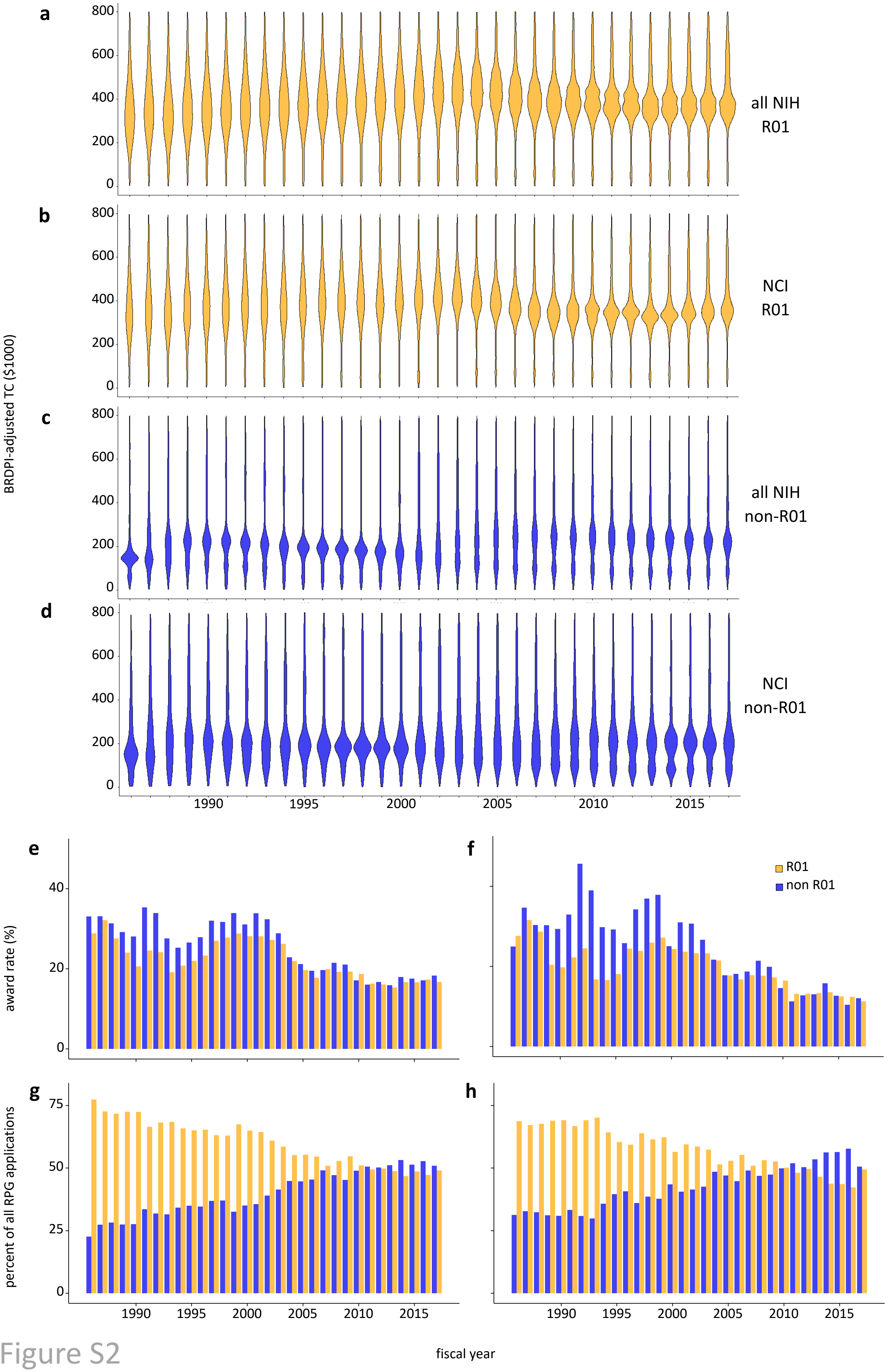

a
b
c
d
e
f
g
h
BRDPI-adjusted TC ($1000)
0
200
400
600
800
all NIH
R01
NCI
R01
all NIH
non-R01
NCI
non-R01
1990
1995
2000
2005
2010
2015
award rate (%)
0
20
40
R01
non R01
percent of all RPG applications
0
25
50
75
fiscal year

Figure S2

**Figure S2. Proliferation and expansion of non-R01 mechanisms is sufficient to explain the declining number of R01 awards.** Data on award size, award rate, and proportion of applications received for the NIH and the NCI between fiscal years 1986 and 2017; R01s are shown in orange and all other non-R01 RPGs in blue. Distribution of award size for the entire NIH and for the NCI is shown as total costs (TC) in thousands of dollars for (**a,b**) R01 awards and (**c,d**) all other non-R01 research project grants (RPGs), adjusted for inflation to 2017 dollars using the Biomedical Research and Development Price Index (BRDPI). Across all of NIH, the inflation-adjusted mean budget size has remained unchanged. At NCI, R01 budgets have not detectably changed, and non-R01 budgets have declined approximately $6000 each year (see Supplemental Materials for details). For (**e**) all of NIH and (**f**) the NCI, award rates over the same time frame for R01 (orange) and non-R01 (blue) applications. Also, for (**g**) NIH and (**h**) NCI over the same time frame, R01 (orange) and non-R01 (blue) applications as a proportion of all grant applications.

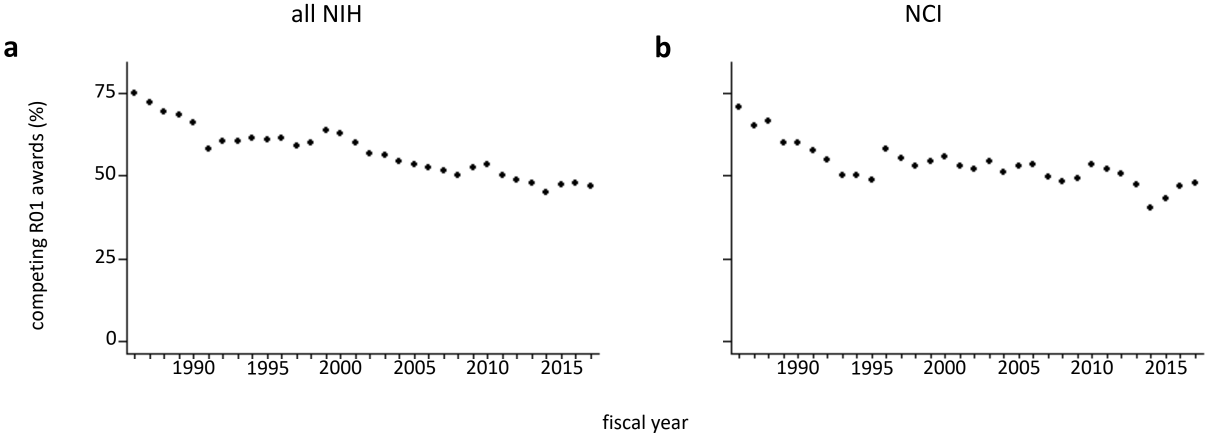

**Figure S3. R01s as a relative fraction of the NIH portfolio.** For (**a**) NIH and (**b**) NCI, the percentage of competing T1 and T2 R01 awards as a fraction of all research grant awards, in fiscal years 1986 to 2017.

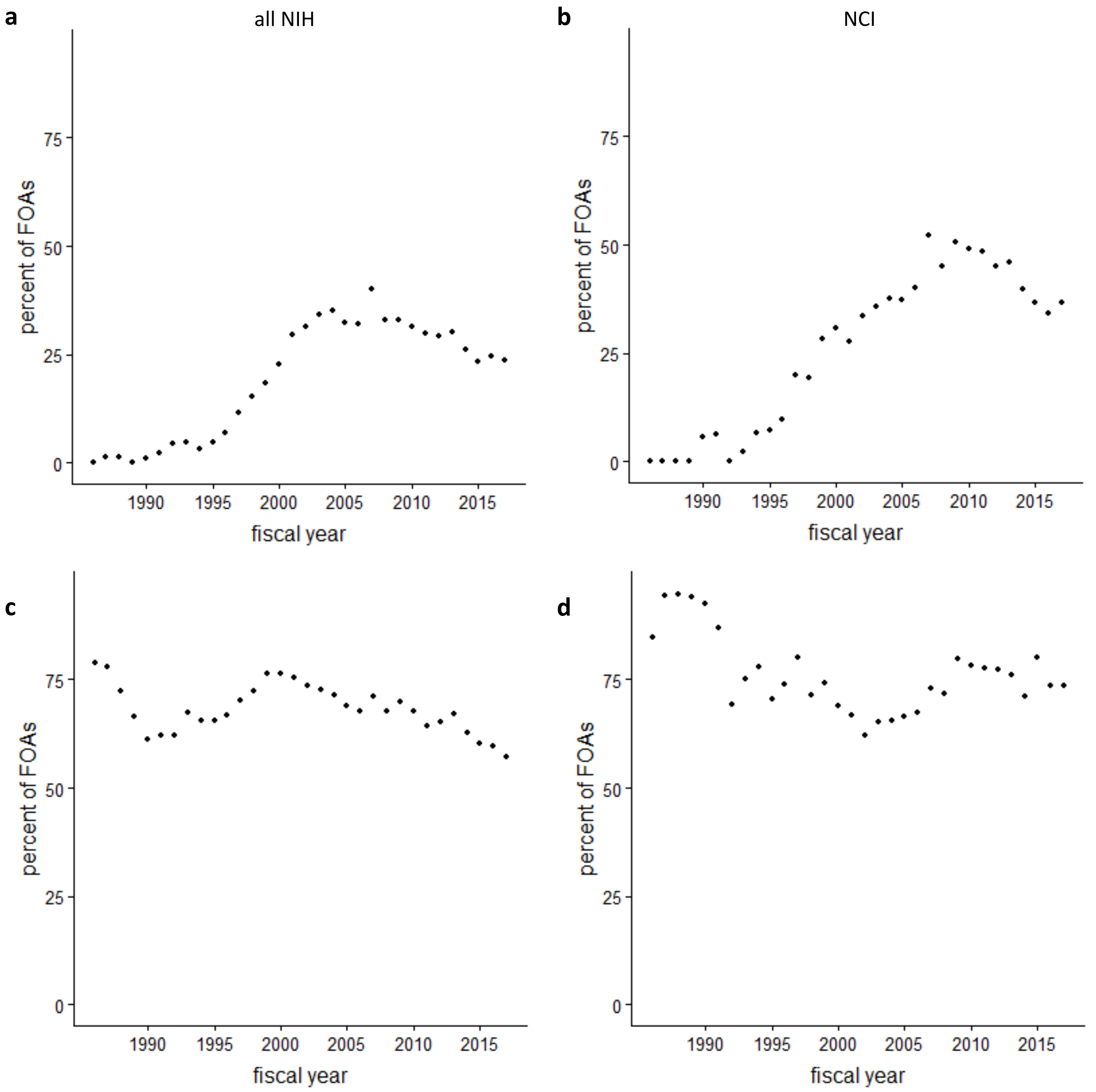

**Figure S4. A small number of mechanisms account for a large percentage of funding opportunity announcements (FOAs) issued annually by NIH and NCI.** (**a, b**) Percent of FOAs issued annually by NIH (left) and NCI (right) through the R21 mechanism. (**c, d**) Percent of FOAs issued annually by NIH (left) and NCI (right) through the R01, R21, R03, and U01 mechanisms combined.

Figure S4

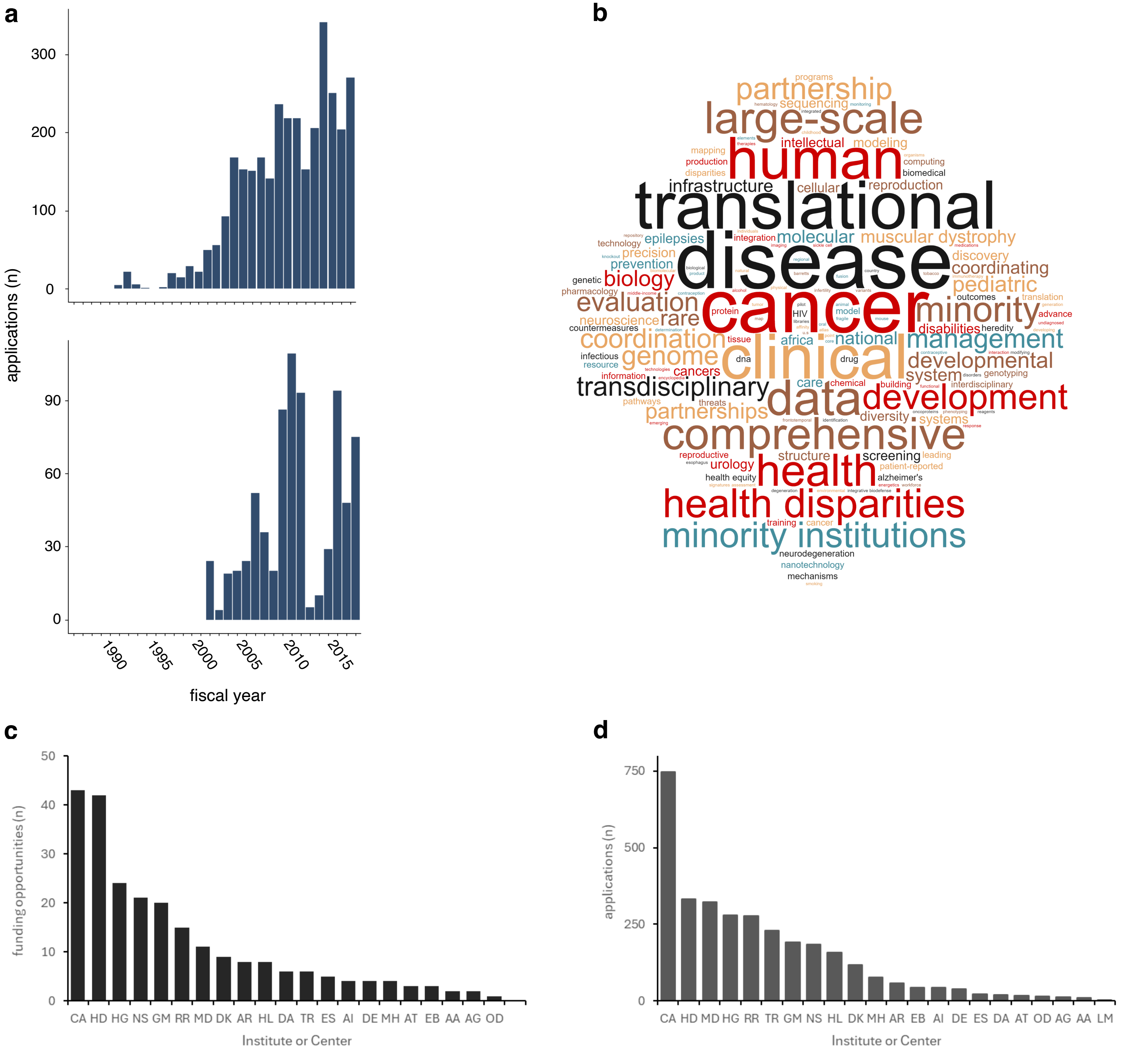

**Figure S5. Analysis of U54 usage.** (**a**) Number of U54 applications to NIH (top) or NCI (bottom) by year. (**b**) Themes of U54 Funding Opportunity Announcements. Word size correlates with frequency of appearance in the titles of U54 funding opportunity announcements, with larger words being more common and smaller words being less common. (**c**) Number of funding opportunities using the U54 across all years by Institute or Center (IC). (**d**) Number of U54 applications across all years, by IC.

Figure S5

Table S1. NIH-wide data on mechanism use.

| Mechanism | First yr | Most yr | Last yr | First Count | Most Count | Last Count | Most:First | Last:First | Duration | active 2017 | >30 appls | >6 yrs |
|---|---|---|---|---|---|---|---|---|---|---|---|---|
| R21 | 1986 | 2016 | 2017 | 4 | 16139 | 15257 | 4034.8 | 3814.3 | 31 | 1 | 1 | 1 |
| R34 | 2003 | 2011 | 2017 | 1 | 717 | 585 | 717 | 585 | 14 | 1 | 1 | 1 |
| U24 | 1992 | 2017 | 2017 | 1 | 301 | 301 | 301 | 301 | 25 | 1 | 1 | 1 |
| R24 | 1989 | 2013 | 2017 | 1 | 586 | 209 | 586 | 209 | 28 | 1 | 1 | 1 |
| U19 | 1993 | 2014 | 2017 | 1 | 272 | 142 | 272 | 142 | 24 | 1 | 1 | 1 |
| P20 | 1986 | 2004 | 2017 | 2 | 352 | 163 | 176 | 81.5 | 31 | 1 | 1 | 1 |
| UC4 | 2011 | 2017 | 2017 | 1 | 69 | 69 | 69 | 69 | 6 | 1 | 1 | 1 |
| U54 | 1991 | 2014 | 2017 | 4 | 341 | 270 | 85.3 | 67.5 | 26 | 1 | 1 | 1 |
| U44 | 1994 | 2010 | 2017 | 1 | 39 | 33 | 39 | 33 | 23 | 1 | 1 | 1 |
| R25 | 1986 | 2011 | 2017 | 15 | 563 | 483 | 37.5 | 32.2 | 31 | 1 | 1 | 1 |
| UM1 | 2011 | 2014 | 2017 | 2 | 157 | 48 | 78.5 | 24 | 6 | 1 | 1 | 1 |
| R35 | 1986 | 2016 | 2017 | 51 | 958 | 861 | 18.8 | 16.9 | 31 | 1 | 1 | 1 |
| R33 | 1999 | 2017 | 2017 | 19 | 318 | 318 | 16.7 | 16.7 | 18 | 1 | 1 | 1 |
| DP3 | 2009 | 2012 | 2017 | 5 | 180 | 57 | 36 | 11.4 | 8 | 1 | 1 | 1 |
| P41 | 1986 | 2010 | 2017 | 2 | 98 | 22 | 49 | 11 | 31 | 1 | 1 | 1 |
| S06 | 1988 | 2006 | 2017 | 1 | 46 | 11 | 46 | 11 | 29 | 1 | 1 | 1 |
| R44 | 1986 | 2016 | 2017 | 183 | 1905 | 1589 | 10.4 | 8.7 | 31 | 1 | 1 | 1 |
| D43 | 1988 | 2011 | 2017 | 8 | 119 | 53 | 14.9 | 6.6 | 29 | 1 | 1 | 1 |
| R42 | 1996 | 2016 | 2017 | 29 | 265 | 184 | 9.1 | 6.3 | 21 | 1 | 1 | 1 |
| P42 | 1987 | 1995 | 2017 | 4 | 42 | 25 | 10.5 | 6.3 | 30 | 1 | 1 | 1 |
| R15 | 1986 | 2017 | 2017 | 327 | 1812 | 1812 | 5.5 | 5.5 | 31 | 1 | 1 | 1 |
| U01 | 1986 | 2015 | 2017 | 410 | 1964 | 1791 | 4.8 | 4.4 | 31 | 1 | 1 | 1 |
| R03 | 1986 | 2004 | 2017 | 675 | 3796 | 2788 | 5.6 | 4.1 | 31 | 1 | 1 | 1 |
| R41 | 1994 | 2016 | 2017 | 289 | 1511 | 1119 | 5.2 | 3.9 | 23 | 1 | 1 | 1 |
| R56 | 2005 | 2008 | 2017 | 111 | 420 | 376 | 3.8 | 3.4 | 12 | 1 | 1 | 1 |
| SC3 | 2007 | 2008 | 2017 | 25 | 103 | 79 | 4.1 | 3.2 | 10 | 1 | 1 | 1 |
| P30 | 1986 | 2011 | 2017 | 36 | 220 | 108 | 6.1 | 3 | 31 | 1 | 1 | 1 |
| G08 | 1986 | 1995 | 2017 | 22 | 210 | 56 | 9.5 | 2.5 | 31 | 1 | 1 | 1 |
| R36 | 2004 | 2014 | 2017 | 16 | 52 | 39 | 3.3 | 2.4 | 13 | 1 | 1 | 1 |
| UH2 | 2009 | 2014 | 2017 | 41 | 239 | 93 | 5.8 | 2.3 | 8 | 1 | 1 | 1 |
| R43 | 1986 | 2004 | 2017 | 1676 | 5346 | 3719 | 3.2 | 2.2 | 31 | 1 | 1 | 1 |
| U18 | 1990 | 2012 | 2017 | 5 | 49 | 10 | 9.8 | 2 | 27 | 1 | 1 | 1 |
| R01 | 1986 | 2017 | 2017 | 18537 | 34205 | 34205 | 1.8 | 1.8 | 31 | 1 | 1 | 1 |
| U34 | 2009 | 2015 | 2017 | 10 | 56 | 18 | 5.6 | 1.8 | 8 | 1 | 1 | 1 |
| SC2 | 2007 | 2008 | 2017 | 47 | 130 | 76 | 2.8 | 1.6 | 10 | 1 | 1 | 1 |
| P50 | 1986 | 2003 | 2017 | 100 | 261 | 156 | 2.6 | 1.6 | 31 | 1 | 1 | 1 |
| P01 | 1986 | 2004 | 2017 | 336 | 571 | 311 | 1.7 | 0.9 | 31 | 1 | 1 | 1 |
| S10 | 1991 | 2010 | 2017 | 406 | 1612 | 366 | 4 | 0.9 | 26 | 1 | 1 | 1 |
| DP1 | 2004 | 2008 | 2017 | 239 | 515 | 208 | 2.2 | 0.9 | 13 | 1 | 1 | 1 |
| SC1 | 2007 | 2008 | 2017 | 90 | 216 | 64 | 2.4 | 0.7 | 10 | 1 | 1 | 1 |
| R18 | 1986 | 1991 | 2017 | 61 | 239 | 34 | 3.9 | 0.6 | 31 | 1 | 1 | 1 |
| DP5 | 2011 | 2011 | 2017 | 108 | 108 | 44 | 1 | 0.4 | 6 | 1 | 1 | 1 |
| U56 | 2001 | 2004 | 2017 | 8 | 45 | 2 | 5.6 | 0.3 | 16 | 1 | 1 | 1 |
| DP2 | 2007 | 2007 | 2017 | 2181 | 2181 | 499 | 1 | 0.2 | 10 | 1 | 1 | 1 |
| U43 | 1993 | 2012 | 2017 | 10 | 51 | 2 | 5.1 | 0.2 | 24 | 1 | 1 | 1 |
| R37 | 1986 | 1987 | 2017 | 177 | 260 | 34 | 1.5 | 0.2 | 31 | 1 | 1 | 1 |
| U10 | 1986 | 1994 | 2017 | 110 | 252 | 1 | 2.3 | 0 | 31 | 1 | 1 | 1 |
| RF1 | 2012 | 2017 | 2017 | 7 | 134 | 134 | 19.1 | 19.1 | 5 | 1 | 1 | 0 |
| UG1 | 2014 | 2016 | 2017 | 53 | 95 | 85 | 1.8 | 1.6 | 3 | 1 | 1 | 0 |
| U2C | 2014 | 2016 | 2017 | 1 | 42 | 14 | 42 | 14 | 3 | 1 | 1 | 0 |
| UG3 | 2015 | 2017 | 2017 | 3 | 257 | 257 | 85.7 | 85.7 | 2 | 1 | 1 | 0 |
| R50 | 2016 | 2016 | 2017 | 218 | 218 | 83 | 1 | 0.4 | 1 | 1 | 1 | 0 |
| R61 | 2016 | 2017 | 2017 | 103 | 147 | 147 | 1.4 | 1.4 | 1 | 1 | 1 | 0 |
| SB1 | 2016 | 2017 | 2017 | 72 | 104 | 104 | 1.4 | 1.4 | 1 | 1 | 1 | 0 |
| OT2 | 2016 | 2017 | 2017 | 20 | 41 | 41 | 2.1 | 2.1 | 1 | 1 | 1 | 0 |
| P60 | 1986 | 2012 | 2016 | 7 | 60 | 4 | 8.6 | 0.6 | 30 | 0 | 1 | 1 |
| R55 | 1991 | 1991 | 2013 | 305 | 305 | 1 | 1 | 0 | 22 | 0 | 1 | 1 |

| | | | | | | | | | | | | |
|---|---|---|---|---|---|---|---|---|---|---|---|---|
| R29 | 1986 | 1995 | 2000 | 58 | 2516 | 1 | 43.4 | 0 | 14 | 0 | 1 | 1 |
| S07 | 1991 | 1991 | 2003 | 628 | 628 | 117 | 1 | 0.2 | 12 | 0 | 1 | 1 |
| RC1 | 2000 | 2008 | 2010 | 40 | 86 | 1 | 2.2 | 0 | 10 | 0 | 1 | 1 |
| UC1 | 2000 | 2004 | 2005 | 8 | 130 | 93 | 16.3 | 11.6 | 5 | 0 | 1 | 0 |
| R22 | 1986 | 1986 | 1989 | 96 | 96 | 64 | 1 | 0.7 | 3 | 0 | 1 | 0 |
| DP7 | 2013 | 2013 | 2014 | 105 | 105 | 66 | 1 | 0.6 | 1 | 0 | 1 | 0 |
| R23 | 1986 | 1986 | 1987 | 1075 | 1075 | 298 | 1 | 0.3 | 1 | 0 | 1 | 0 |
| RL1 | 2007 | 2007 | 2007 | 51 | 51 | 51 | 1 | 1 | 0 | 0 | 1 | 0 |
| PN1 | 2004 | 2004 | 2004 | 81 | 81 | 81 | 1 | 1 | 0 | 0 | 1 | 0 |
| U42 | 1991 | 2006 | 2017 | 2 | 23 | 5 | 11.5 | 2.5 | 26 | 1 | 0 | 1 |
| P40 | 1991 | 2003 | 2017 | 10 | 16 | 2 | 1.6 | 0.2 | 26 | 1 | 0 | 1 |
| P51 | 1991 | 2008 | 2017 | 1 | 3 | 2 | 3 | 2 | 26 | 1 | 0 | 1 |
| U41 | 1992 | 2012 | 2017 | 1 | 21 | 14 | 21 | 14 | 25 | 1 | 0 | 1 |
| S21 | 2001 | 2011 | 2017 | 6 | 12 | 3 | 2 | 0.5 | 16 | 1 | 0 | 1 |
| U2R | 2003 | 2004 | 2017 | 1 | 24 | 11 | 24 | 11 | 14 | 1 | 0 | 1 |
| R90 | 2004 | 2004 | 2017 | 17 | 17 | 4 | 1 | 0.2 | 13 | 1 | 0 | 1 |
| UL1 | 2006 | 2007 | 2017 | 12 | 21 | 11 | 1.8 | 0.9 | 11 | 1 | 0 | 1 |
| UH3 | 2013 | 2016 | 2017 | 1 | 18 | 12 | 18 | 12 | 4 | 1 | 0 | 0 |
| UF1 | 2013 | 2016 | 2017 | 3 | 3 | 1 | 1 | 0.3 | 4 | 1 | 0 | 0 |
| P2C | 2014 | 2015 | 2017 | 6 | 26 | 6 | 4.3 | 1 | 3 | 1 | 0 | 0 |
| UM2 | 2014 | 2017 | 2017 | 1 | 3 | 3 | 3 | 3 | 3 | 1 | 0 | 0 |
| RM1 | 2015 | 2016 | 2017 | 7 | 17 | 5 | 2.4 | 0.7 | 2 | 1 | 0 | 0 |
| OT3 | 2016 | 2017 | 2017 | 5 | 16 | 16 | 3.2 | 3.2 | 1 | 1 | 0 | 0 |
| RC2 | 2017 | 2017 | 2017 | 11 | 11 | 11 | 1 | 1 | 0 | 1 | 0 | 0 |
| G12 | 1991 | 2003 | 2014 | 8 | 9 | 7 | 1.1 | 0.9 | 23 | 0 | 0 | 1 |
| M01 | 1991 | 1993 | 2008 | 19 | 24 | 7 | 1.3 | 0.4 | 17 | 0 | 0 | 1 |
| S11 | 1994 | 2004 | 2011 | 1 | 14 | 2 | 14 | 2 | 17 | 0 | 0 | 1 |
| UC7 | 2006 | 2016 | 2016 | 2 | 2 | 2 | 1 | 1 | 10 | 0 | 0 | 1 |
| UH1 | 1996 | 2003 | 2004 | 5 | 5 | 3 | 1 | 0.6 | 8 | 0 | 0 | 1 |
| RL5 | 2007 | 2014 | 2014 | 3 | 21 | 21 | 7 | 7 | 7 | 0 | 0 | 1 |
| S22 | 2001 | 2001 | 2007 | 8 | 8 | 2 | 1 | 0.3 | 6 | 0 | 0 | 1 |
| PN2 | 2005 | 2005 | 2010 | 20 | 20 | 6 | 1 | 0.3 | 5 | 0 | 0 | 0 |
| UA5 | 2011 | 2011 | 2015 | 1 | 1 | 1 | 1 | 1 | 4 | 0 | 0 | 0 |
| UG4 | 2016 | 2016 | 2016 | 11 | 11 | 11 | 1 | 1 | 0 | 0 | 0 | 0 |
| PL1 | 2007 | 2007 | 2007 | 11 | 11 | 11 | 1 | 1 | 0 | 0 | 0 | 0 |
| RC4 | 2010 | 2010 | 2010 | 6 | 6 | 6 | 1 | 1 | 0 | 0 | 0 | 0 |
| RL9 | 2007 | 2007 | 2007 | 5 | 5 | 5 | 1 | 1 | 0 | 0 | 0 | 0 |
| RC3 | 2010 | 2010 | 2010 | 2 | 2 | 2 | 1 | 1 | 0 | 0 | 0 | 0 |
| R28 | 2013 | 2013 | 2013 | 1 | 1 | 1 | 1 | 1 | 0 | 0 | 0 | 0 |
| RL2 | 2007 | 2007 | 2007 | 1 | 1 | 1 | 1 | 1 | 0 | 0 | 0 | 0 |

Table S2. Data on mechanism use at NCI.

| Mechanism | First yr | Most yr | Last yr | First Count | Most Count | Last Count | Most:First | Last:First | Duration | active 2017 | >30 appls | >6 yrs |
|---|---|---|---|---|---|---|---|---|---|---|---|---|
| R21 | 1986 | 2017 | 2017 | 4 | 13214 | 13214 | 3303.5 | 3303.5 | 31 | 1 | 1 | 1 |
| R34 | 2003 | 2011 | 2017 | 1 | 717 | 585 | 717 | 585 | 14 | 1 | 1 | 1 |
| R24 | 1989 | 2013 | 2017 | 1 | 585 | 209 | 585 | 209 | 28 | 1 | 1 | 1 |
| R25 | 1987 | 2015 | 2017 | 1 | 496 | 422 | 496 | 422 | 30 | 1 | 1 | 1 |
| U19 | 1993 | 2014 | 2017 | 1 | 271 | 141 | 271 | 141 | 24 | 1 | 1 | 1 |
| U24 | 1992 | 2017 | 2017 | 1 | 217 | 217 | 217 | 217 | 25 | 1 | 1 | 1 |
| P20 | 1989 | 2004 | 2017 | 2 | 352 | 79 | 176 | 39.5 | 28 | 1 | 1 | 1 |
| R35 | 1988 | 2016 | 2017 | 6 | 777 | 716 | 129.5 | 119.3 | 29 | 1 | 1 | 1 |
| U54 | 1991 | 2014 | 2017 | 4 | 312 | 195 | 78 | 48.8 | 26 | 1 | 1 | 1 |
| UC4 | 2011 | 2017 | 2017 | 1 | 69 | 69 | 69 | 69 | 6 | 1 | 1 | 1 |
| UM1 | 2011 | 2013 | 2017 | 2 | 105 | 47 | 52.5 | 23.5 | 6 | 1 | 1 | 1 |
| P41 | 1986 | 2010 | 2017 | 2 | 98 | 22 | 49 | 11 | 31 | 1 | 1 | 1 |
| S06 | 1988 | 2006 | 2017 | 1 | 46 | 11 | 46 | 11 | 29 | 1 | 1 | 1 |
| DP3 | 2009 | 2012 | 2017 | 5 | 180 | 57 | 36 | 11.4 | 8 | 1 | 1 | 1 |
| D43 | 1988 | 2011 | 2017 | 8 | 119 | 53 | 14.9 | 6.6 | 29 | 1 | 1 | 1 |
| U10 | 1987 | 2011 | 2017 | 17 | 208 | 1 | 12.2 | 0.1 | 30 | 1 | 1 | 1 |
| R44 | 1986 | 2016 | 2017 | 145 | 1556 | 1281 | 10.7 | 8.8 | 31 | 1 | 1 | 1 |
| P42 | 1987 | 1995 | 2017 | 4 | 42 | 25 | 10.5 | 6.3 | 30 | 1 | 1 | 1 |
| U18 | 1990 | 2012 | 2017 | 5 | 49 | 10 | 9.8 | 2 | 27 | 1 | 1 | 1 |
| G08 | 1986 | 1995 | 2017 | 22 | 210 | 56 | 9.5 | 2.5 | 31 | 1 | 1 | 1 |
| R33 | 2001 | 2006 | 2017 | 11 | 105 | 10 | 9.5 | 0.9 | 16 | 1 | 1 | 1 |
| P30 | 1986 | 2011 | 2017 | 22 | 205 | 98 | 9.3 | 4.5 | 31 | 1 | 1 | 1 |
| R42 | 1996 | 2016 | 2017 | 26 | 221 | 151 | 8.5 | 5.8 | 21 | 1 | 1 | 1 |
| R18 | 1986 | 1991 | 2017 | 35 | 236 | 34 | 6.7 | 1 | 31 | 1 | 1 | 1 |
| R03 | 1986 | 2004 | 2017 | 513 | 3428 | 1973 | 6.7 | 3.8 | 31 | 1 | 1 | 1 |
| U44 | 2002 | 2010 | 2017 | 6 | 39 | 33 | 6.5 | 5.5 | 15 | 1 | 1 | 1 |
| R15 | 1986 | 2015 | 2017 | 279 | 1576 | 1527 | 5.6 | 5.5 | 31 | 1 | 1 | 1 |
| U34 | 2009 | 2015 | 2017 | 10 | 56 | 18 | 5.6 | 1.8 | 8 | 1 | 1 | 1 |
| R41 | 1994 | 2016 | 2017 | 235 | 1191 | 865 | 5.1 | 3.7 | 23 | 1 | 1 | 1 |
| U01 | 1986 | 2015 | 2017 | 319 | 1408 | 1070 | 4.4 | 3.4 | 31 | 1 | 1 | 1 |
| SC3 | 2007 | 2008 | 2017 | 25 | 103 | 79 | 4.1 | 3.2 | 10 | 1 | 1 | 1 |
| UH2 | 2009 | 2015 | 2017 | 38 | 154 | 75 | 4.1 | 2 | 8 | 1 | 1 | 1 |
| S10 | 1991 | 2010 | 2017 | 406 | 1612 | 366 | 4 | 0.9 | 26 | 1 | 1 | 1 |
| R56 | 2005 | 2008 | 2017 | 110 | 384 | 370 | 3.5 | 3.4 | 12 | 1 | 1 | 1 |
| R43 | 1986 | 2004 | 2017 | 1267 | 4291 | 3022 | 3.4 | 2.4 | 31 | 1 | 1 | 1 |
| R36 | 2004 | 2014 | 2017 | 16 | 52 | 39 | 3.3 | 2.4 | 13 | 1 | 1 | 1 |
| SC2 | 2007 | 2008 | 2017 | 47 | 124 | 76 | 2.6 | 1.6 | 10 | 1 | 1 | 1 |
| SC1 | 2007 | 2008 | 2017 | 89 | 210 | 59 | 2.4 | 0.7 | 10 | 1 | 1 | 1 |
| P50 | 1986 | 1999 | 2017 | 99 | 225 | 109 | 2.3 | 1.1 | 31 | 1 | 1 | 1 |
| DP1 | 2004 | 2008 | 2017 | 239 | 515 | 206 | 2.2 | 0.9 | 13 | 1 | 1 | 1 |
| R01 | 1986 | 2017 | 2017 | 15872 | 28145 | 28145 | 1.8 | 1.8 | 31 | 1 | 1 | 1 |
| P01 | 1986 | 2004 | 2017 | 263 | 440 | 216 | 1.7 | 0.8 | 31 | 1 | 1 | 1 |
| R37 | 1986 | 1987 | 2017 | 150 | 223 | 34 | 1.5 | 0.2 | 31 | 1 | 1 | 1 |
| DP5 | 2011 | 2011 | 2017 | 108 | 108 | 44 | 1 | 0.4 | 6 | 1 | 1 | 1 |
| DP2 | 2007 | 2007 | 2017 | 2181 | 2181 | 493 | 1 | 0.2 | 10 | 1 | 1 | 1 |
| UG3 | 2015 | 2017 | 2017 | 3 | 245 | 245 | 81.7 | 81.7 | 2 | 1 | 1 | 0 |
| RF1 | 2012 | 2017 | 2017 | 7 | 134 | 134 | 19.1 | 19.1 | 5 | 1 | 1 | 0 |
| OT2 | 2016 | 2017 | 2017 | 20 | 41 | 41 | 2.1 | 2.1 | 1 | 1 | 1 | 0 |
| U2C | 2015 | 2016 | 2017 | 23 | 42 | 14 | 1.8 | 0.6 | 2 | 1 | 1 | 0 |
| UG1 | 2015 | 2016 | 2017 | 54 | 95 | 85 | 1.8 | 1.6 | 2 | 1 | 1 | 0 |
| SB1 | 2016 | 2017 | 2017 | 72 | 104 | 104 | 1.4 | 1.4 | 1 | 1 | 1 | 0 |
| R61 | 2016 | 2017 | 2017 | 103 | 147 | 147 | 1.4 | 1.4 | 1 | 1 | 1 | 0 |
| R29 | 1986 | 1995 | 2000 | 47 | 2172 | 1 | 46.2 | 0 | 14 | 0 | 1 | 1 |
| P60 | 1986 | 2012 | 2016 | 7 | 60 | 4 | 8.6 | 0.6 | 30 | 0 | 1 | 1 |
| RC1 | 2000 | 2008 | 2010 | 40 | 86 | 1 | 2.2 | 0 | 10 | 0 | 1 | 1 |
| R55 | 1991 | 1991 | 2013 | 250 | 250 | 1 | 1 | 0 | 22 | 0 | 1 | 1 |
| S07 | 1991 | 1991 | 2003 | 628 | 628 | 117 | 1 | 0.2 | 12 | 0 | 1 | 1 |

| | | | | | | | | | | | | |
|---|---|---|---|---|---|---|---|---|---|---|---|---|
| UC1 | 2000 | 2004 | 2005 | 8 | 130 | 93 | 16.3 | 11.6 | 5 | 0 | 1 | 0 |
| DP7 | 2013 | 2013 | 2014 | 105 | 105 | 66 | 1 | 0.6 | 1 | 0 | 1 | 0 |
| RL1 | 2007 | 2007 | 2007 | 47 | 47 | 47 | 1 | 1 | 0 | 0 | 1 | 0 |
| PN1 | 2004 | 2004 | 2004 | 81 | 81 | 81 | 1 | 1 | 0 | 0 | 1 | 0 |
| R22 | 1986 | 1986 | 1989 | 96 | 96 | 64 | 1 | 0.7 | 3 | 0 | 1 | 0 |
| R23 | 1986 | 1986 | 1987 | 940 | 940 | 265 | 1 | 0.3 | 1 | 0 | 1 | 0 |
| U2R | 2003 | 2004 | 2017 | 1 | 24 | 11 | 24 | 11 | 14 | 1 | 0 | 1 |
| U41 | 1992 | 2012 | 2017 | 1 | 21 | 14 | 21 | 14 | 25 | 1 | 0 | 1 |
| U42 | 1991 | 2006 | 2017 | 2 | 23 | 5 | 11.5 | 2.5 | 26 | 1 | 0 | 1 |
| P51 | 1991 | 2008 | 2017 | 1 | 3 | 2 | 3 | 2 | 26 | 1 | 0 | 1 |
| S21 | 2001 | 2011 | 2017 | 6 | 12 | 3 | 2 | 0.5 | 16 | 1 | 0 | 1 |
| UL1 | 2006 | 2007 | 2017 | 12 | 21 | 11 | 1.8 | 0.9 | 11 | 1 | 0 | 1 |
| P40 | 1991 | 2003 | 2017 | 10 | 16 | 2 | 1.6 | 0.2 | 26 | 1 | 0 | 1 |
| U56 | 2003 | 2003 | 2017 | 12 | 12 | 2 | 1 | 0.2 | 14 | 1 | 0 | 1 |
| R90 | 2004 | 2004 | 2017 | 17 | 17 | 4 | 1 | 0.2 | 13 | 1 | 0 | 1 |
| UH3 | 2013 | 2016 | 2017 | 1 | 15 | 9 | 15 | 9 | 4 | 1 | 0 | 0 |
| P2C | 2014 | 2015 | 2017 | 6 | 26 | 6 | 4.3 | 1 | 3 | 1 | 0 | 0 |
| OT3 | 2016 | 2017 | 2017 | 5 | 16 | 16 | 3.2 | 3.2 | 1 | 1 | 0 | 0 |
| UM2 | 2014 | 2017 | 2017 | 1 | 3 | 3 | 3 | 3 | 3 | 1 | 0 | 0 |
| RM1 | 2015 | 2016 | 2017 | 7 | 17 | 5 | 2.4 | 0.7 | 2 | 1 | 0 | 0 |
| UF1 | 2013 | 2016 | 2017 | 3 | 3 | 1 | 1 | 0.3 | 4 | 1 | 0 | 0 |
| RC2 | 2017 | 2017 | 2017 | 11 | 11 | 11 | 1 | 1 | 0 | 1 | 0 | 0 |
| U43 | 2017 | 2017 | 2017 | 1 | 1 | 1 | 1 | 1 | 0 | 1 | 0 | 0 |
| S11 | 1994 | 2004 | 2011 | 1 | 14 | 2 | 14 | 2 | 17 | 0 | 0 | 1 |
| RL5 | 2007 | 2014 | 2014 | 2 | 21 | 21 | 10.5 | 10.5 | 7 | 0 | 0 | 1 |
| M01 | 1991 | 1993 | 2008 | 19 | 24 | 7 | 1.3 | 0.4 | 17 | 0 | 0 | 1 |
| G12 | 1991 | 2003 | 2014 | 8 | 9 | 7 | 1.1 | 0.9 | 23 | 0 | 0 | 1 |
| UC7 | 2006 | 2016 | 2016 | 2 | 2 | 2 | 1 | 1 | 10 | 0 | 0 | 1 |
| UH1 | 1996 | 2003 | 2004 | 5 | 5 | 3 | 1 | 0.6 | 8 | 0 | 0 | 1 |
| S22 | 2001 | 2001 | 2007 | 8 | 8 | 2 | 1 | 0.3 | 6 | 0 | 0 | 1 |
| PN2 | 2005 | 2005 | 2010 | 20 | 20 | 6 | 1 | 0.3 | 5 | 0 | 0 | 0 |
| UG4 | 2016 | 2016 | 2016 | 11 | 11 | 11 | 1 | 1 | 0 | 0 | 0 | 0 |
| UA5 | 2015 | 2015 | 2015 | 1 | 1 | 1 | 1 | 1 | 0 | 0 | 0 | 0 |
| R28 | 2013 | 2013 | 2013 | 1 | 1 | 1 | 1 | 1 | 0 | 0 | 0 | 0 |
| RC4 | 2010 | 2010 | 2010 | 6 | 6 | 6 | 1 | 1 | 0 | 0 | 0 | 0 |
| RC3 | 2010 | 2010 | 2010 | 2 | 2 | 2 | 1 | 1 | 0 | 0 | 0 | 0 |
| PL1 | 2007 | 2007 | 2007 | 10 | 10 | 10 | 1 | 1 | 0 | 0 | 0 | 0 |
| RL9 | 2007 | 2007 | 2007 | 4 | 4 | 4 | 1 | 1 | 0 | 0 | 0 | 0 |
| RL2 | 2007 | 2007 | 2007 | 1 | 1 | 1 | 1 | 1 | 0 | 0 | 0 | 0 |
